# Mouse lymph node colonization models predict human tumor metastasis in melanoma

**DOI:** 10.64898/2026.09.22.753675

**Authors:** Brendan K. Ball, Reem Al-Humadi, Sabrina Zdravkovic, Ilayda Ilerten, Brooks A. Benard, Brooke E. Howitt, Andrew J. Gentles

## Abstract

Aggressive tumor progression and distant metastases of cancers such as cutaneous melanoma are thought to be preceded by regional lymph node (LN) infiltration. Previous mouse studies have selectively enriched for tumor cells with enhanced LN migratory capacity but the mechanistic biology underpinning this proclivity has not been directly linked to human outcomes. Likewise, the exact role LN colonization plays in distant metastasis in humans remains unclear. To translate and conserve findings from mouse models to humans, we used a computational framework termed Translatable Components Regression (TransComp-R) to integrate mouse LN tumor samples with human primary and metastatic tumors. We identified transcriptional programs of early- and late-stage mouse tumor generations that stratified primary and metastatic tumors from patients with melanoma. These transcriptional programs associated with immune function, cell cycle checkpoints, and DNA-to-protein biosynthesis pathways. We then employed EcoTyper and identified six distinct melanoma-specific cellular communities (ecotypes) comprising co-occurring cell states. Of these six melanoma ecotypes, we identified a community strongly associated with primary tumor cells, while the other five represented a spectrum of metastatic tumor microenvironments with increased adverse outcomes. These ecotypes and their significantly expressed genes may contribute to the tumor-immune microenvironment that facilitates LN colonization in melanoma. We further validated the physical co-localization of these melanoma ecotypes using spatial transcriptomics. Our integration of pre-clinical models with human data reveals potential biological pathways and transcriptomic signatures for future investigation and supports the utility of mouse models in studying cancer metastasis.

**GRAPHICAL ABSTRACT:** 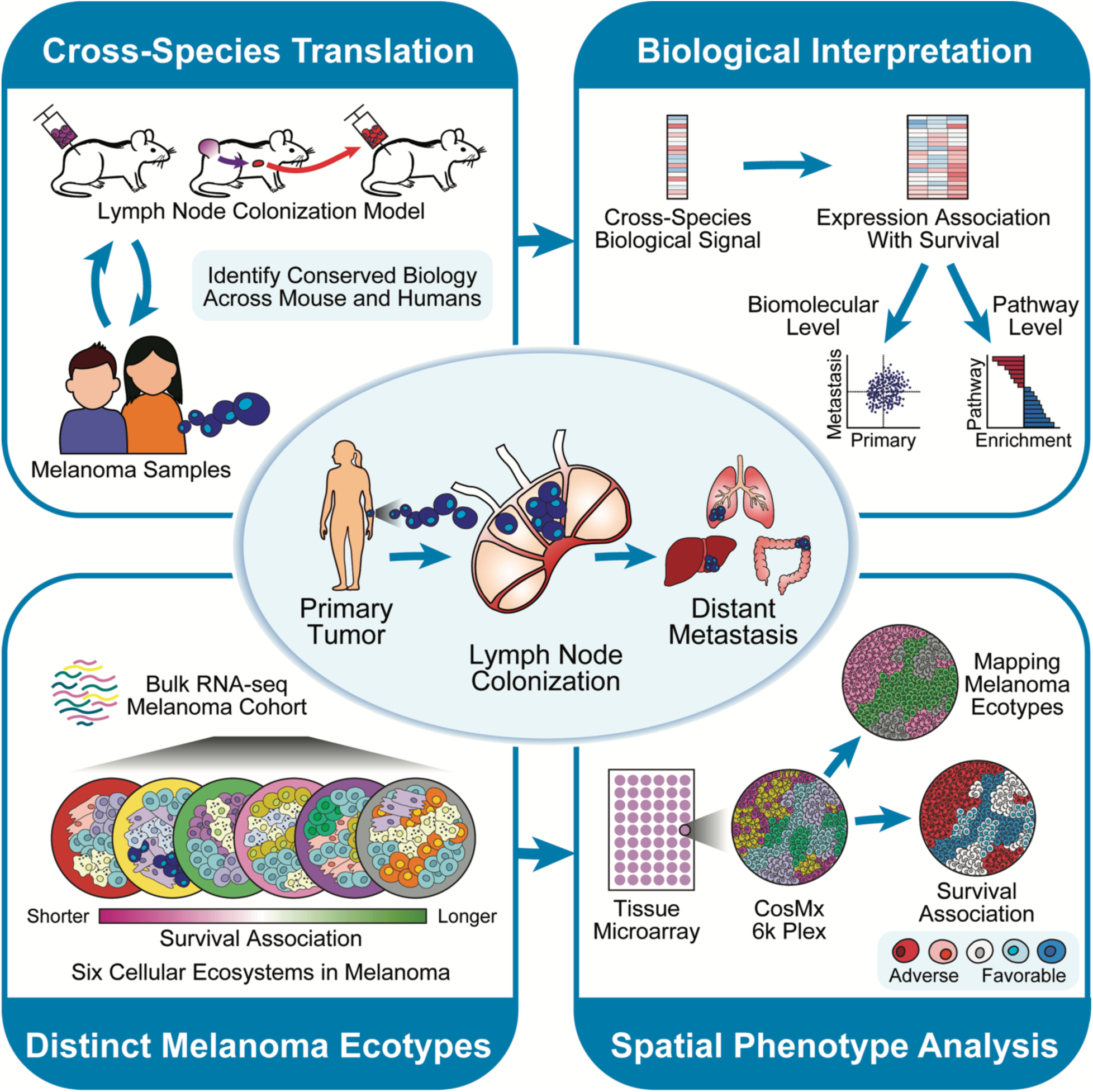

## INTRODUCTION

Infiltration of the lymph node (LN) by malignant cells is a pivotal indication of metastasis and adverse outcomes in many cancers^1^. LNs are essential organs of the lymphatic and immune system responsible for filtering out harmful substances and abnormal cells as well as acting as educational hubs for the immune system^2,3^. In cancers such as melanoma, LN involvement is thought to precede distant metastasis by inducing and remodeling tumor-immune tolerance, which is subsequently exported systemically^4–6^. This alteration of the LNs may be attributed to factors such as stromal reprogramming by cancer cells^7^, modulation of the LN immune microenvironment^8,9^, in addition to structural and morphological changes to high endothelial venules^10,11^. Despite ongoing efforts to understand this biological process, the exact mechanism by which primary tumors become metastatic is unclear.

Cross-species analyses offer a path to identify conserved biological signals linked to tumor metastasis and outcomes. For example, pre-clinical studies in mouse models often support the use of more invasive experiments and controlled longitudinal studies but may not fully emulate physiological processes in humans^12,13^. This lack of generalization of pre-clinical findings to humans often leads to translational failure in the clinic^14,15^. Alternatively, clinical studies can directly support advancements in our understanding of cancer pathology, but often face challenges of heterogeneity in human populations^16^. We hypothesized that synergizing information between mouse and human transcriptomic data could reveal conserved LN-associated biomarkers predictive of metastatic outcomes in melanoma and support the relevance of these mouse models to human disease.

We leveraged a computational framework called Translatable Components Regression (TransComp-R)^17–20^ to integrate publicly available pre-clinical and human transcriptomics data. This approach enables the projection of tumor samples from human melanoma onto mouse-collected tumor data, allowing us to identify transcriptomic signatures prognostic of metastatic and primary outcomes in humans. Simultaneously, we apply a series of downstream computational modeling techniques to further interpret these transcriptomic signatures and identify DNA-to-protein synthesis along with cell checkpoint pathways enriched in metastatic samples. We then applied EcoTyper, a framework to systematically identify cellular states and communities (ecotypes) in melanoma. As a result, we identified six distinct ecotypes unique to melanoma and further demonstrate their spatial co-localization using spatial transcriptomics. Together, this multi-scale and systems approach reveals biological signals in mice predictive of tumor metastasis in humans along with distinct melanoma ecotypes with varying survival outcomes.

## RESULTS

### Mouse lymph node colonization models stratify human primary and metastatic melanoma outcomes

To understand if we could identify conserved biological signals indicative of cancer metastasis across species, we combined mouse and human data using the TransComp-R framework. We tested this question by synthesizing two separate datasets: a mouse LN colonization model with sequential tumor generations increasing in selectivity to the LN (GSE117529)^6^, and a clinically annotated primary and metastatic melanoma human dataset (GSE46517)^21^ (**Figure 1a**).

**Figure 1.**
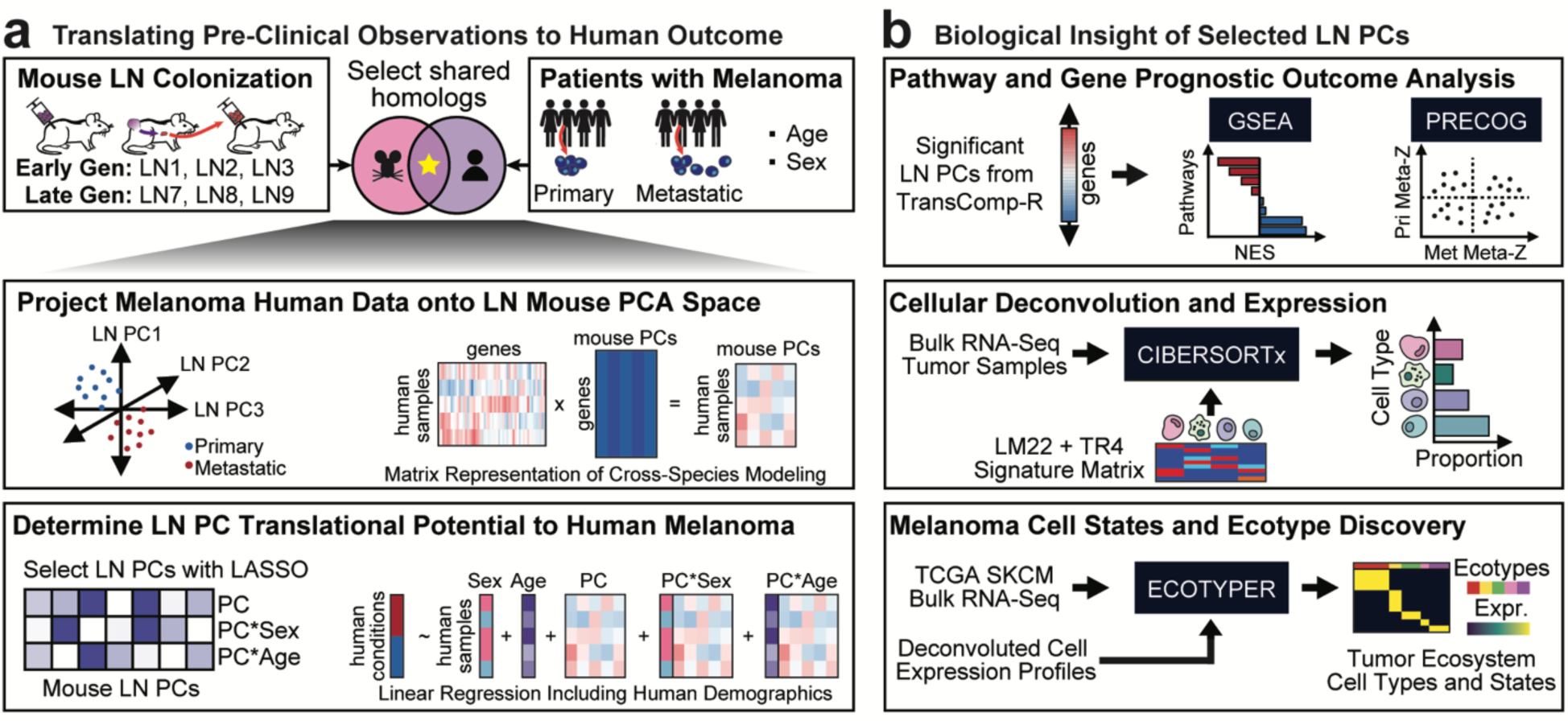
Cross-species modeling and downstream biological interpretation. **(a)** Mouse LN colonization models are synthesized with primary and metastatic tumor samples from patients diagnosed with melanoma. Mouse LN PCs combined with human demographic variables are regressed against tumor metastasis outcomes to determine PC predictability. **(b)** The LN PCs determined to have significant predictive ability in distinguishing primary from metastatic tumors are interpreted for biological relevance through pathway enrichment analysis, identification of genes with favorable or adverse outcomes, cell type deconvolution, and discovery of melanoma-specific ecosystems with EcoTyper.

We first defined a mouse LN principal component analysis (PCA) space based on bulk RNA-seq from passaged tumors and then projected the human melanoma samples into this space. In order to identify conserved transcriptional programs associated with human metastasis, we employed LASSO (Least Absolute Shrinkage and Selection Operator)^22^, a variable feature selection method, to regress the projected mouse LN principal components (PCs) against metastasis outcomes while incorporating interaction terms for potential confounders of sex and age. To interpret the biological variance encoded in the significant PCs, we perform pathway enrichment analysis and cell type deconvolution (**Figure 1b**). We then identify distinct cell type and state ecosystems relevant to melanoma. Our approach focuses on identifying both pathway-level and specific biomolecular changes that drive tumor metastasis.

After matching for 10,436 shared gene homologs between the mouse and human data (**Supplementary Figure S1**), we evaluated the nine mouse LN PCs that explained a cumulative 81.3% variance in mouse for their ability to predict metastasis outcomes in human melanoma. Of the nine mouse LN PCs, we identified six PCs that significantly distinguished between human primary and metastatic tumor outcomes when selected for a combined model by LASSO (**Figure 2a**).

**Figure 2.**
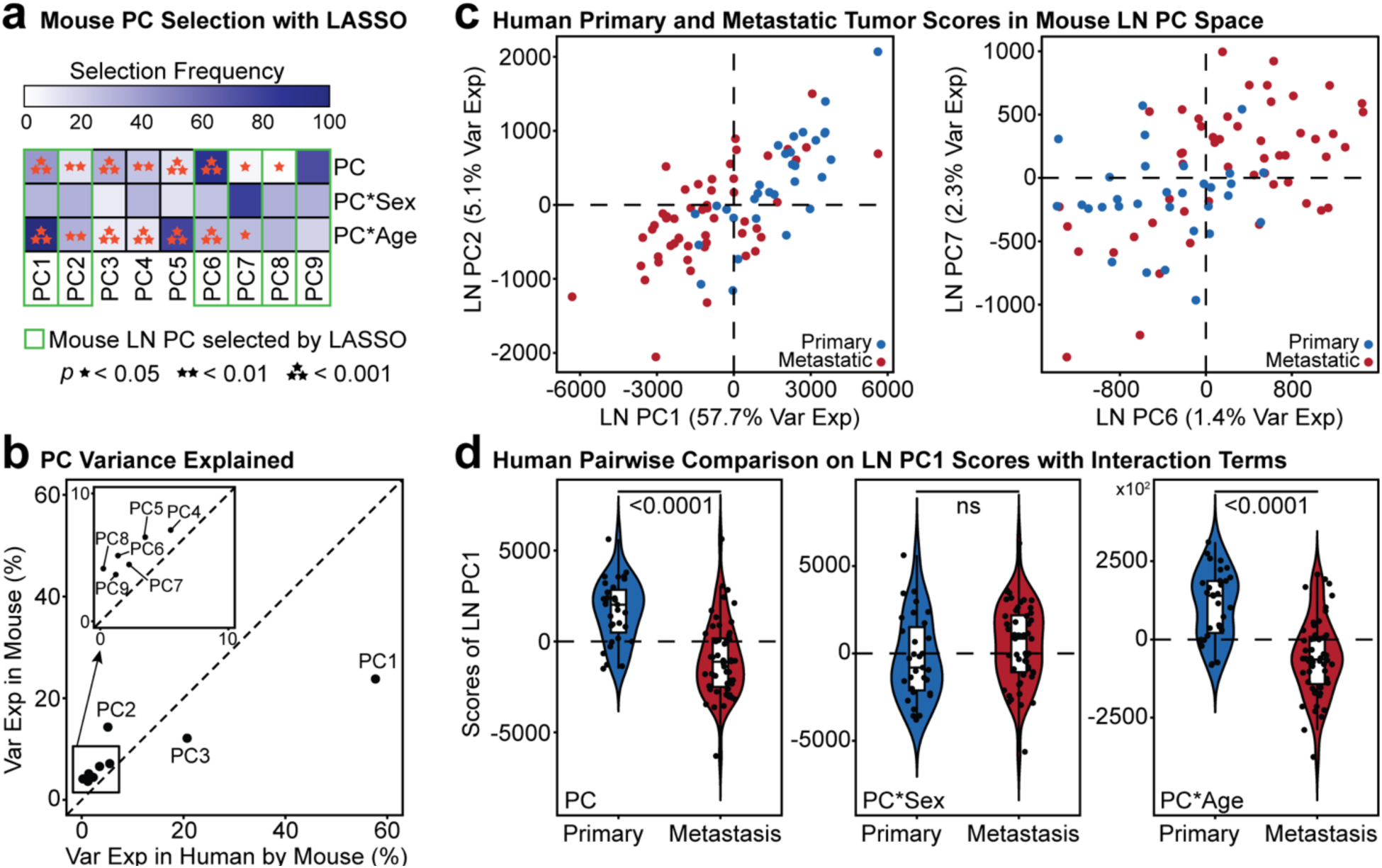
Mouse LN PC1 separates human primary and metastatic melanoma samples. **(a)** Mouse LN PC feature selection with the LASSO model. Selection frequency is the number of times the LASSO model selects the specific mouse LN PC at random. The model p values were determined by a generalized linear regression. **(b)** Comparison of the percent variance explained by the mouse PCs and its translational ability to explain human variance. **(c)** Projected TransComp-R plots of the human samples into mouse LN PC spaces. **(d)** Mann-Whitney pairwise comparison of the selected LN PC1 across interaction terms of sex and age, with Benjamini-Hochberg-adjusted p values.

We evaluated the six significant mouse LN PCs characterized by their ability to explain the overall variance in the human melanoma dataset. Of the six mouse LN PCs, mouse LN PC1 had the highest proportion of variance explained in the human population by mouse models (57.7%), which indicates potential for this mouse LN PC to be informative in the human melanoma context (**Figure 2b**). We visualized the human projections onto the mouse PC spaces and demonstrated distinct separation across the metastatic and primary human tumor samples (**Figure 2c**). Although other mouse LN PCs such as PC6 and PC7 portrayed separation of primary and metastatic tumor samples of humans, they explained less than 10% of the variance in the human gene expression data (**Supplementary Figure S1**).

We also sought to determine if demographic variables of sex and age contributed to the ability to distinguish metastatic and primary outcomes. While we found that the LN PC1 scores with sex interaction effects were not significant, we found interaction between age and LN PC1 was statistically significant (p<0.0001), indicating that associations between LN PC1 and metastasis are also contributed by patient age (**Figure 2d**).

### Gene signatures regulating immune function, protein biosynthesis, and cell cycle checkpoint pathways associate with cancer prognosis and lymph node metastasis

Having established that mouse LN PC1 could distinguish between human primary and metastatic tumor samples, we next sought to identify the biological pathways underlying this signature that are conserved between human and mouse. We performed Gene Set Enrichment Analysis (GSEA) with the KEGG (Kyoto Encyclopedia of Genes and Genomes) and Hallmark curations to identify overrepresented pathways. For KEGG, we found pathways involved in cell cycle, mismatch repair, DNA replication, RNA degradation, and metabolic pathways such as purine metabolism to be enriched in metastatic tumors, whereas immune signaling such as cytokine-cytokine receptor interaction enrichment was associated with primary tumors (**Figure 3a**). Among the enriched Hallmark pathways, we identified programs associated with the cell cycle checkpoints, E2F targets, MYC targets, DNA repair, and oxidative phosphorylation as upregulated among metastatic tumors (**Figure 3b**). Among the differences between primary and metastatic tumor pathway enrichment, we found an overrepresentation of genes contributing to the p53 pathway, downregulation of KRAS signaling, and interferon immune signaling in primary tumors associated with primary tumors, whereas pathways associated with cellular proliferation and transcriptional mechanisms were more enriched in metastatic tumors. This difference may indicate the biological shift from immune pathway upregulation in early cancer stages to cellular proliferation pathways in later cancer stages.

**Figure 3.**
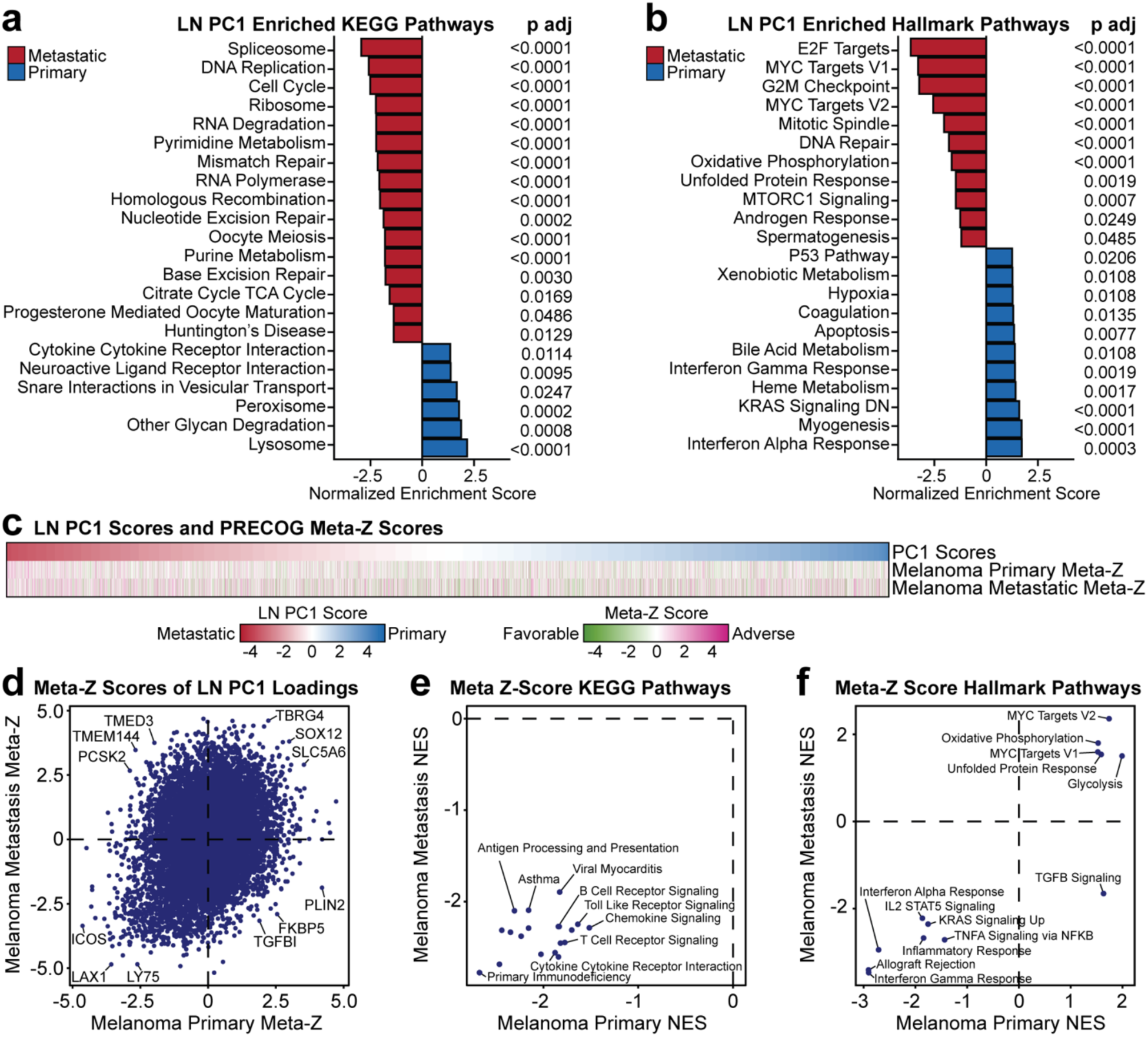
Biological pathway enrichment analysis and computational prognosis analysis. **(a)** Enriched pathways under the KEGG database. **(b)** Enriched pathways under the Hallmark database. **(c)** Genes and their scores on PC1 labeled by their primary and metastatic melanoma meta-Z scores from PRECOG. **(d)** Comparison of the PRECOG meta-Z scores for both primary and metastatic outcomes and LN PC1 scores. **(e)** Shared KEGG pathways pre-ranked on genes and their meta-Z scores in primary and metastatic melanoma. **(f)** Shared Hallmark pathways pre-ranked on genes and their meta-Z scores in primary and metastatic melanoma.

To interrogate the prognostic associations of the genes defining mouse LN PC1, we used our previously described PRECOG (Prediction of Clinical Outcomes from Genomics)^23,24^ resource to identify genes with favorable or adverse cancer outcomes in primary and metastatic melanoma (**Figure 3c**). We compared the primary and metastatic melanoma meta-Z scores to identify genes associated with favorable or adverse clinical outcomes (**Figure 3d**). Some genes (*ICOS*, *LAX1*, and *LY75*) associated with completely favorable outcomes whereas others (*TBRG4*, *SOX12*, and *SLC5A6*) associated with completely adverse outcomes in both primary and metastatic melanoma signatures. Meanwhile, genes such as *TMED3*, *TMEM144*, and *PCSK2* were favorable under primary but adverse under metastatic conditions. Genes such as *PLIN2*, *FKBP5*, and *TGFBI* were associated with adverse outcomes in primary and favorable outcomes in metastatic melanoma. For instance, reduced expression of *FKBP5* in cancer triggers hyperphosphorylation of Akt and decreased cell death^25^ and has been shown to promote or suppress tumor growth in melanoma^26,27^. These genes may inform active immune responses consistent with favorable outcomes in both primary and metastatic melanoma, and tumor-intrinsic activity such as metabolism and proliferation with adverse outcomes in both primary and metastatic melanoma. Alternatively, genes with inverse clinical outcomes across primary and metastatic melanoma samples may represent a transition from a primary site to eventual metastasis.

We next re-ranked the genes contributing to mouse LN PC1 by their respective PRECOG meta-Z scores and ran GSEA to determine if certain biological pathways contained direct or indirect relationships with respect to their cancer prognosis. After GSEA, we first selected biological pathways that associate with outcomes in both primary and metastatic melanoma (**Supplementary Figure S2**). The entire list of enriched pathways is available in **Supplementary Figure S3**. From the KEGG pathway database, we identified 18 shared pathways, all of which were involved in favorable outcomes in both primary and metastatic melanoma (**Figure 3e**). These enriched pathways were primarily associated with immune signaling. Within the Hallmark curation, we matched 13 shared pathways (**Figure 3f**). Of these, all pathways were assigned with favorable or adverse outcomes in both primary and metastatic melanoma except for one. Of the shared enriched pathways, TGFβ (Transforming growth factor β) signaling was associated with adverse outcomes in primary melanoma, but favorable outcomes in metastatic melanoma. The TGFβ pathway regulates cell growth and differentiation, tissue homeostasis, and immune function. This is particularly interesting because it is the only enriched pathway shared between the primary and metastatic tumor that has an inverse association, while the dual favorable and adverse nature may indicate a transitional shift across biological signaling pathways associated with TGFβ during tumor metastasis.

### Identification of cell-type specific transcriptional differences between primary and metastatic melanoma samples

Cell-type specific gene expression can be estimated from large bulk RNA-seq datasets using deconvolution and digital cytometry with CIBERSORTx (Cell-type Identification by Estimating Relative Subsets of RNA Transcripts)^28^. We used 408 samples of bulk RNA-seq data containing human primary and metastatic skin cutaneous melanoma (SKCM) tumor samples from The Cancer Genome Atlas (TCGA) and deconvoluted the expression data using CIBERSORTx (**Figure 4a**). We then imputed cell-specific gene expression profiles and filtered the analysis to only include mouse LN PC1 genes driving tumor type separation. This approach enables us to associate mouse LN PC1-specific genes that can predict between primary and metastatic outcomes with the cell types by which they are expressed.

**Figure 4.**
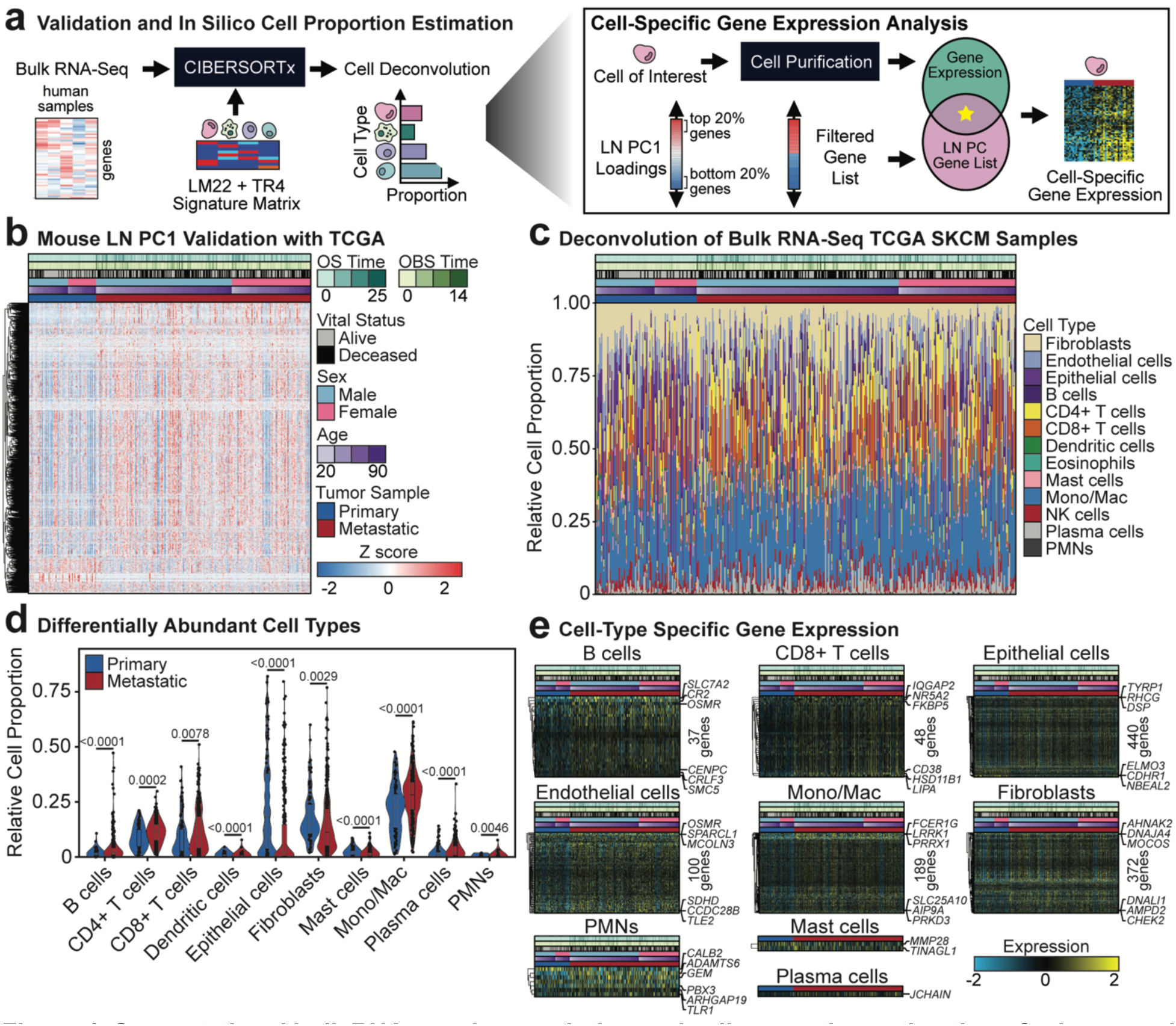
Computational bulk RNA-seq deconvolution and cell proportion estimation of primary and metastatic melanoma tumor samples. **(a)** Approach of identifying LN-associated metastasis in cells using the TCGA SKCM dataset. **(b)** Significant genes filtered by the top and bottom 20% scores from mouse LN PC1 (Mann-Whitney pairwise test, Benjamini-Hochberg adjusted false discovery rate (FDR)<0.05). The overall survival and observed survival time are represented in years **(c)** Estimated cell proportions across all tumor samples. **(d)** Pairwise comparison to identify differentially abundant cell types. (Mann-Whitney pairwise test, Benjamini-Hochberg adjusted FDR<0.05) **(e)** Cell-specific gene expression differentially expressed between primary and metastatic tumor samples (Mann-Whitney pairwise test, Benjamini-Hochberg adjusted FDR<0.05). The top and bottom three (upregulated and downregulated) genes on the heatmap are labeled.

We first filtered genes from the bulk RNA-seq dataset to shared genes within the LN PC1, and of these, 1,778 genes were differentially expressed between the primary and metastatic tumors from the TransComp-R model (**Figure 4b**). From our bulk RNA-seq data with CIBERSORTx, we estimated the proportion of 10 different immune cell types plus epithelial, endothelial, and fibroblasts in each TCGA SKCM sample (**Figure 4c**). Of the 13 cell types we imputed, 10 were differentially abundant between the primary and metastatic tumor samples (**Figure 4d**). This included B cells (FDR<0.0001), CD4+ T cells (FDR=0.0002), CD8+ T cells (FDR=0.0078), dendritic cells (FDR<0.0001), epithelial cells (FDR<0.0001), fibroblasts (FDR=0.0029), mast cells (FDR<0.0001), monocytes and macrophages (FDR<0.0001), plasma cells (FDR<0.0001), and polymorphonuclear leukocytes (PMNs) (FDR=0.0046).

Irrespective of cell population changes, we also questioned if these cell types contained significant transcriptional changes that could also indicate tumor metastasis. Among the 13 cell types, we identified nine that contained at least one significantly differentially expressed gene between primary and metastatic tumor samples (**Figure 4e**). Epithelial and fibroblasts contained the greatest number of differentially expressed genes, with 440 and 372 genes, respectively. Following these were monocytes and macrophages (189 genes), CD8 T cells (48 genes), endothelial cells (100 genes), B cells (37 genes), PMNS (7 genes), mast cells (2 genes; *MMP28* and *TINAGL1*), and plasma cells (1 gene; *JCHAIN*).

### Discovery of melanoma-specific ecotypes reveals distinct tumor ecosystems associated with primary tumor samples and a spectrum of metastatic tumors

We next sought to identify distinct multicellular tumor ecosystems (ecotypes) to characterize melanoma using the TCGA SKCM bulk RNA-seq data. EcoTyper^29^ uses the cell-type gene expression decomposition from CIBERSORTx to identify coherent patterns of transcription representing cell states, and their co-occurrence across the cohort, which organizes them into what we term melanoma ecotypes (MEs). With this melanoma discovery model, we classified a total of 367 patient samples into each of 6 distinct MEs (**Figure 5a-b**). Interestingly, we identified ME5 as a primary tumor-specific ecosystem.

**Figure 5.**
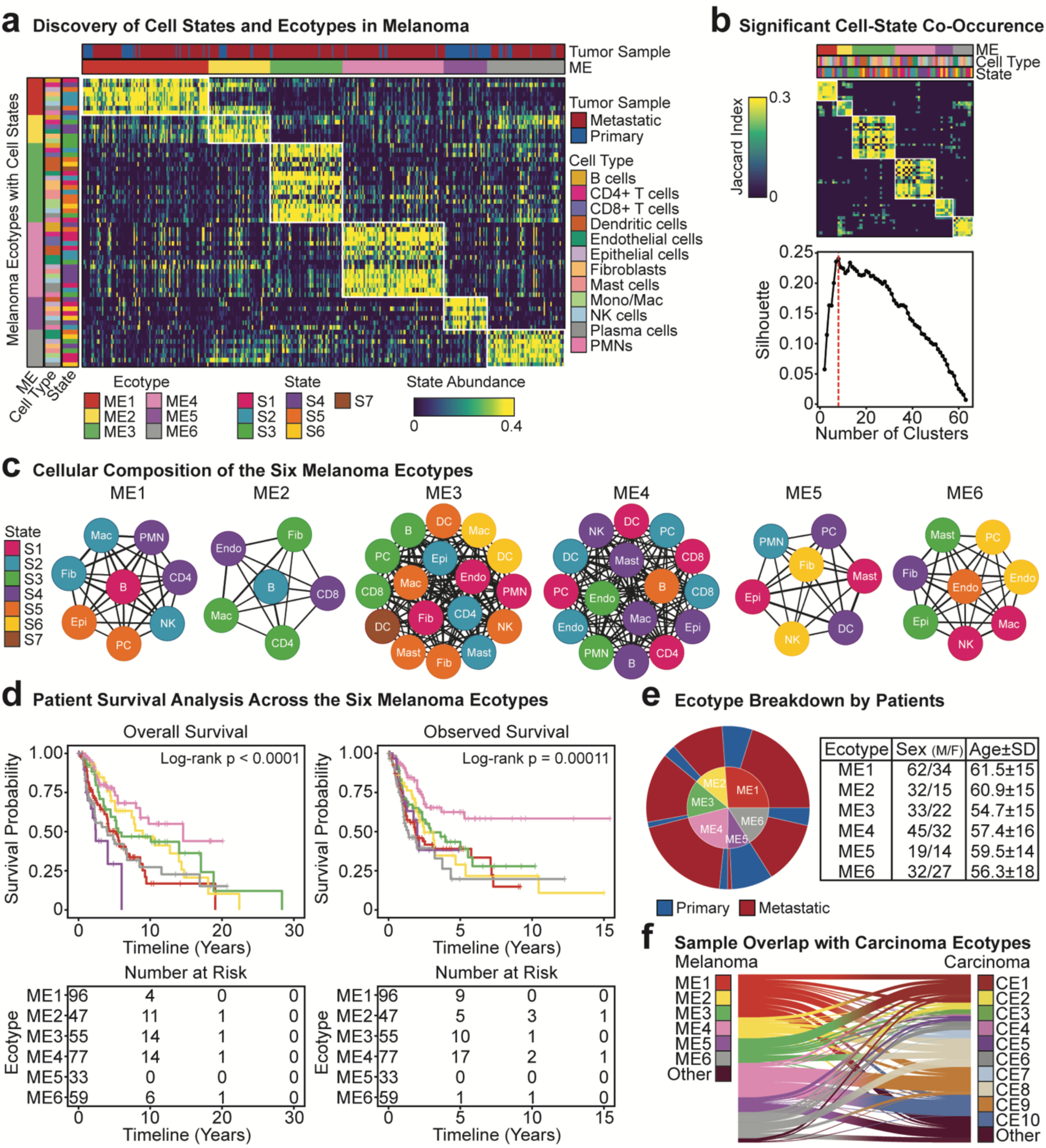
Discovery of melanoma-specific tumor ecosystems. **(a)** Distribution of melanoma ecotypes across the TCGA SKCM tumor samples (n=367 assigned to a ME). **(b)** Significant cell-state co-occurrences and optimized number of clusters (n=8 clusters). **(c)** ME-specific cellular composition and states discovered from EcoTyper. The node and color represent the cell and state, respectively, and the edge thickness is the co-association strength calculated by the Jaccard index. **(d)** Patient overall survival and observed survival across each ME. **(e)** Breakdown of the ME by tumor type, sex, and age distribution. The mean age (in years) for the specific ME and the standard deviation is presented. **(f)** Shared overlap of ecotypes from the melanoma discovery and carcinoma ecotypes (n=318 assigned to a CE).

The six unique MEs were comprised of between 6 to 17 distinct cell states (**Figure 5c**). ME3 contained the largest cellular community (17 distinct cell states across), followed by ME4 (16), ME1 and ME6 (8), ME5 (7), and ME2 (6). We next characterized the survival outcomes by each ME and found ME4 to be associated with the most favorable outcomes under both overall survival (log-ranked p<0.0001) and observed survival (log-ranked p=0.00011) (**Figure 5d**). We also presented the observed survival outcomes, defined as the time interval from sampling to the last follow-up or death, to address the potential out-of-step recording between clinical outcomes and RNA-seq sampling^30^. We confirmed similar demographic distributions across sex and age for all MEs (**Figure 5e**). We also confirmed that these defined tumor ecosystems were not driven by which tissue site the metastatic samples came from in the patients (**Supplementary Figure S4**).

In TCGA, patients who contributed primary tumor samples surprisingly had worse survival outcomes than those contributing metastatic samples. This could arise through biases such as selection of large primaries for genomic profiling that are more likely to have poor outcomes. This observation has been noted by another group addressing a potential out-of-step situation between clinical outcomes and omics data specific to TCGA SKCM^30^. We therefore tested our ME model in a separate bulk RNA-seq dataset (GSE65904) which contained 214 primary and metastatic melanoma samples and validated the MEs (**Supplementary Figure S5**). In this cohort, we found that the patients with tumor samples assigned to an ME4 ecotype had the most favorable tumor ecosystem compared to its other metastatic counterparts. We also observed consistent findings showing that ME5 predominantly contains primary tumor-specific cellular communities. Therefore, our melanoma EcoTyper model identified consistent results, although the survival outcome findings in **Figure 5d** may be specific to TCGA SKCM and GSE65904.

Previously, EcoTyper was used to identify 10 ecotypes in a pan-cancer fashion across 16 solid carcinomas excluding melanoma. To determine if the MEs are similar to these previously established ecotypes, we recovered the carcinoma ecotypes (CEs) in the TCGA SKCM cohort (**Figure 5f**). We traced each patient to their assigned ME from our melanoma-specific model to assigned CEs from the pan-cancer model to compare shared or different tumor ecosystems. Strikingly, ME4, which portended the most favorable survival outcomes in melanoma, overlapped with the favorable survival CE outcomes CE9 and CE10. The CE9 and CE10 tumors were previously reported to be proinflammatory, associated with longer overall survival, and higher immunoreactivity^29^. The other MEs were distributed to other CE assignments. Additional recovery information on the CEs is in **Supplementary Figure S6**.

### Recovery of the TransComp-R predicted human cohort corroborates the discovered primary tumor-specific ecotype

Using our melanoma-specific EcoTyper model, we recovered MEs in the human melanoma data (GSE46517)^21^ we used for TransComp-R. Primary melanoma samples were enriched for ME5 and ME6, whereas ME1-4 were specific to metastatic tumor samples (**Figure 6a,b**). The apparent shift of the ME6 toward primary tumor samples may be an artifact of the limited number of samples specific to the GSE46517 dataset.

**Figure 6.**
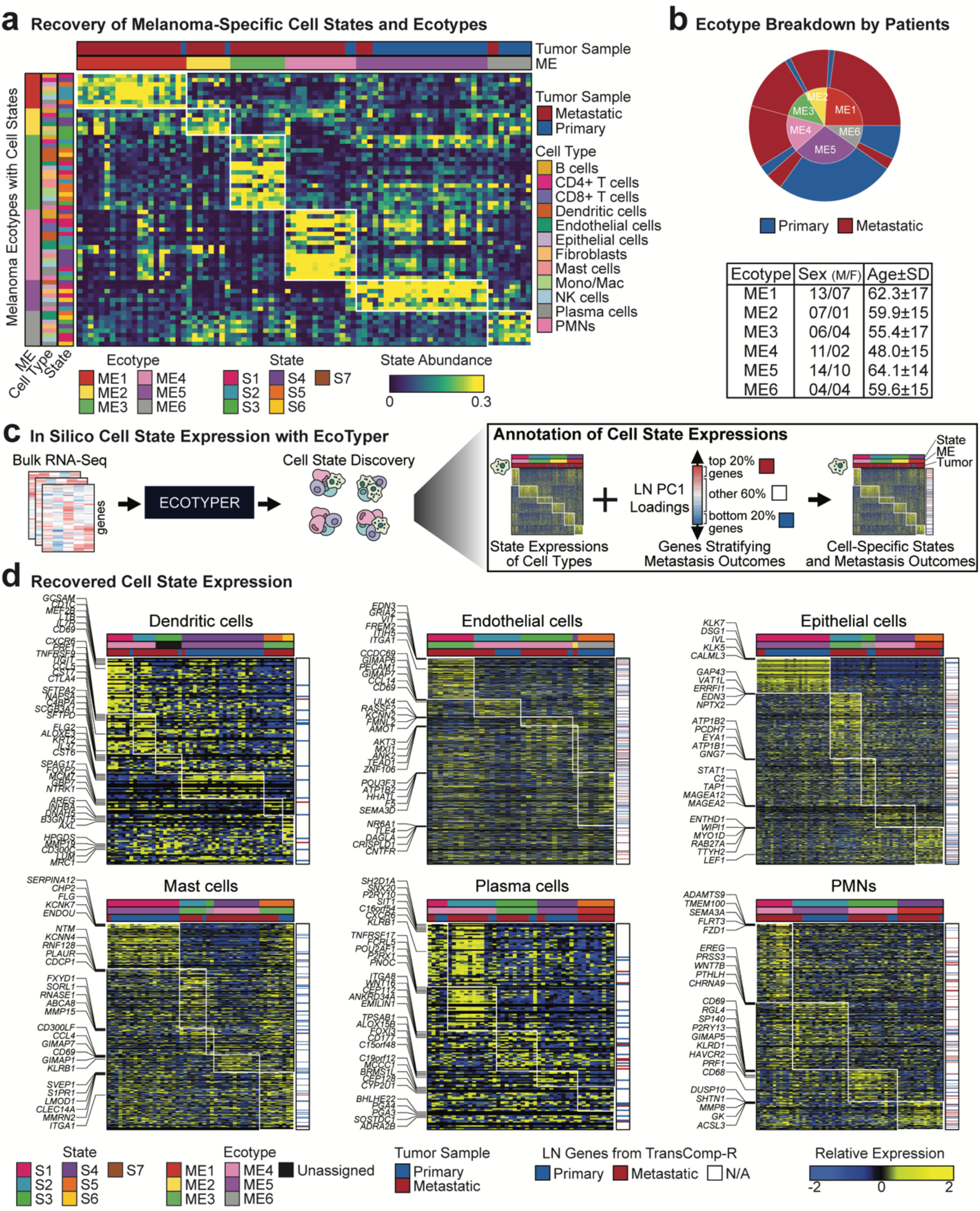
Recovery of melanoma ecotypes. **(a)** Distribution of recovered melanoma ecotypes from the discovered TCGA SKCM EcoTyper model. **(b)** Breakdown of patient demographics by each ME. The age is represented with the mean and standard deviation of the ME. **(c)** Computational workflow of cell state discovery and genes predictive of LN metastasis. **(d)** Recovered cell-state expression annotated by genes predictive of LN metastasis by TransComp-R. These cells contain a cell state unique to primary tumors.

We next mapped the shared mouse-human genes from our TransComp-R model onto EcoTyper cell states to dissect the cell types driving conserved cross-species differences between primary and metastatic melanoma (**Figure 6c**). In particular, we identified six cell types containing a cell-state specific to primary melanoma tumor samples, including dendritic cells (S4), endothelial cells (S5), epithelial cells (S1), mast cells (S1), plasma cells (S4), and PMNs (S2) (**Figure 6d**). Using an independent melanoma cohort, these findings from our recovery model validate the distinct cell-type characteristics of the primary tumor-dominant ME5 identified in our discovery model.

Among the ME5 cell types, we further filtered for genes specific to the primary tumors and found that PMNs contained the greatest number of genes (28 genes) associated with LN metastasis from TransComp-R’s PC1, followed by endothelial cells (27 genes), epithelial cells (16 genes), mast cells (10 genes), dendritic cells (2 genes), and plasma cells (1 gene) (**Supplementary Figure S7**). The other six cell types (B cells, CD4+ T cells, CD8+ T cells, fibroblasts, monocytes and macrophages, and NK cells) and their recovered cell states are available in **Supplementary Figure S8**.

### Biological features and survival characteristics of melanoma ecotypes

After identifying six distinct MEs, we profiled each ME by its clinical survival outcomes, cell-type populations, and enriched genomic features^31^ (**Figure 7a**). Across the six MEs, tumors with elevated abundance of ME1, ME2, ME5, or ME6 were characterized by shorter survival outcomes, whereas ME4 had longer survival outcomes (**Figure 7b**).

**Figure 7.**
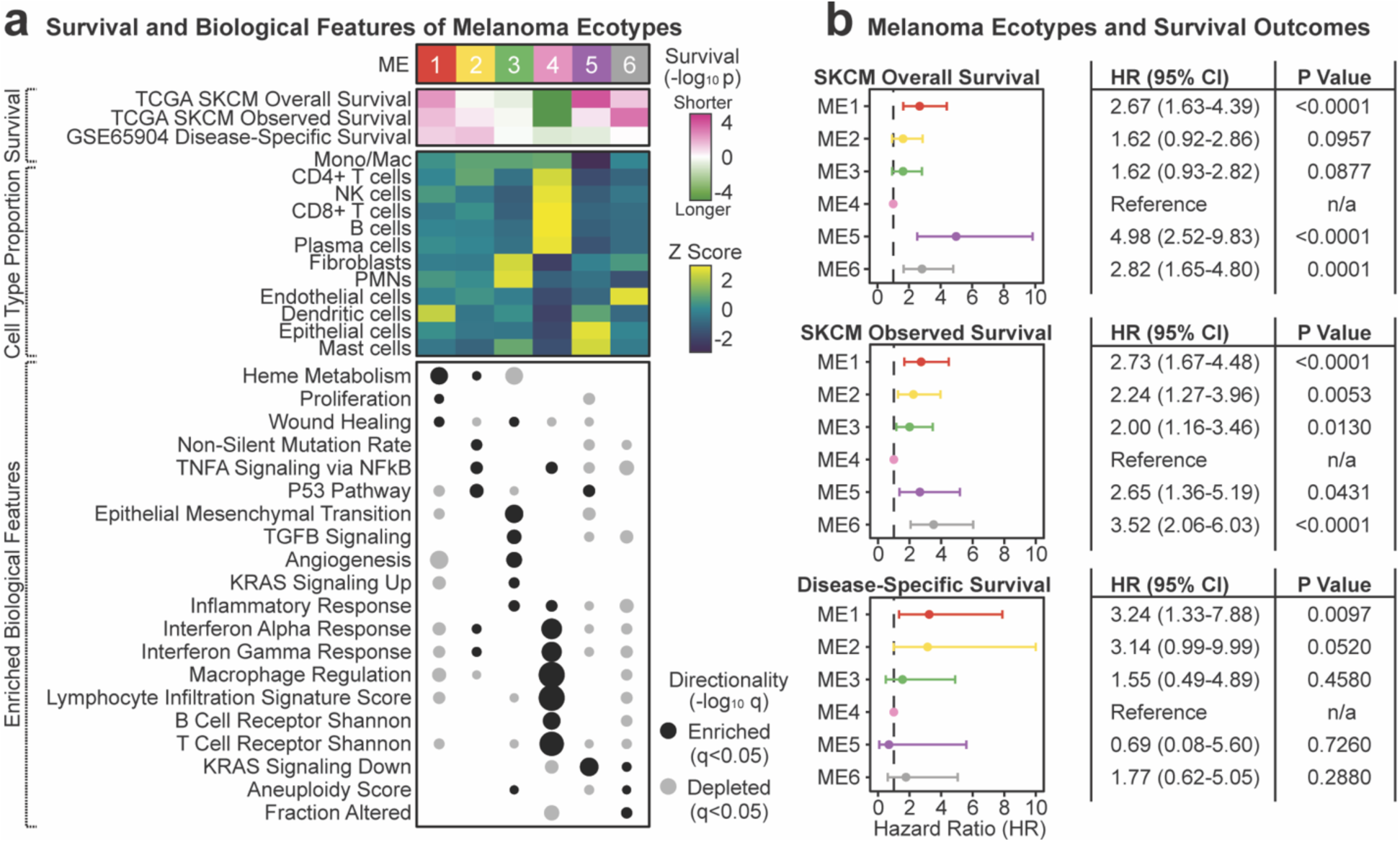
Summary of the melanoma ecotypes. **(a)** Features of the melanoma ecotypes from the TCGA discovery cohort. **Top:** Survival associations across the TCGA SKCM labeled by overall survival and observed survival, as well as the recovery model disease-specific survival from GSE65904. **Middle:** Relative CIBERSORTx cell type proportions represented by Z scores. A greater Z score represents an abundant cell type compared to other MEs. **Bottom:** Enriched GSEA Hallmark and Immunogenomics features grouped by the most abundant ME per tumor. Pairwise testing compared by the specific ME to the other five MEs (FDR q<0.05). Missing entries represent lack of enrichment for the specific ecotype. **(b)** Survival analysis for the SKCM TCGA (overall and observed survival) and GSE65904 (disease-specific survival). Hazard ratios in reference to ME4, the most favorable melanoma ecotype.

Tumors abundant with ME1 are enriched in heme metabolism, proliferation, and wound healing. ME2 is represented by an enrichment for non-silent mutation rate, TNFα signaling via NFκB, and p53 pathway. These pathways suggest that ME2 may associate with mechanisms of tumor suppressor pathways to drive metastasis. Tumors containing ME3 are represented by abundant fibroblasts and PMNs and show biological enrichment for epithelial-to-mesenchymal transition, TGFβ signaling, angiogenesis, KRAS signaling, and inflammatory response. Of all ecotypes, ME4-rich tumors contain a greater abundance of leukocytes and immune cells, explaining ME4’s higher survival odds compared to other cellular communities associated with tumor metastasis. The ME4 communities are characterized by elevated inflammation response, interferon alpha and gamma responses, and immune regulation. The primary tumor-dominant ecotype, ME5, is represented by high abundance of epithelial and mast cells along with enrichment for KRAS signaling suppression. The tumors containing high ME6 are rich in endothelial cells and suppresses KRAS signaling, number of altered chromosome arms (aneuploidy score), and chromosomal instability (fraction altered).

### Melanoma ecotypes spatially co-localize in tissue microarray samples of primary and metastatic samples

To determine if our established MEs are spatially distinct across cellular communities, we performed spatial transcriptomics using the CosMx platform on a tissue microarray (TMA) composed of melanoma samples from 29 individuals. These samples were derived from primary or metastatic melanoma spanning 10 anatomic sites (**Table 1**).

**Table 1.**
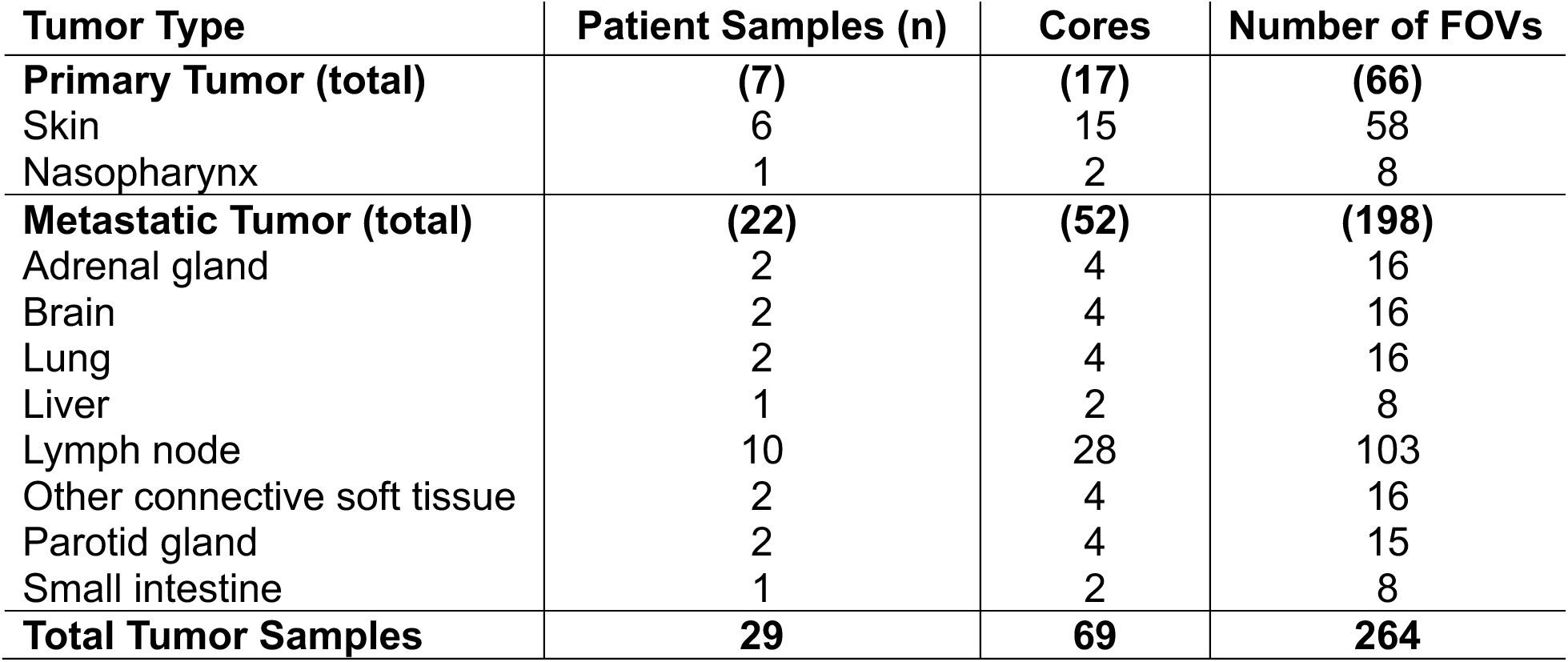
Samples collected from patients with melanoma for spatial transcriptomics.

From the 29 individuals, we imaged 264 fields of view (FOV) from 69 TMA cores of primary and metastatic melanoma, producing a total of 374,263 high-quality cells (**Figure 8a**). These cells comprise 3,276 B cells, 4,131 endothelial cells, 30,094 fibroblasts, 261,674 melanoma cells, 61,367 monocytes and macrophages, 7,149 plasma cells, and 6,572 T cells. The primary samples were largely from skin tissue, whereas the majority of metastasis samples collection locations were from the LNs (**Figure 8b**).

**Figure 8.**
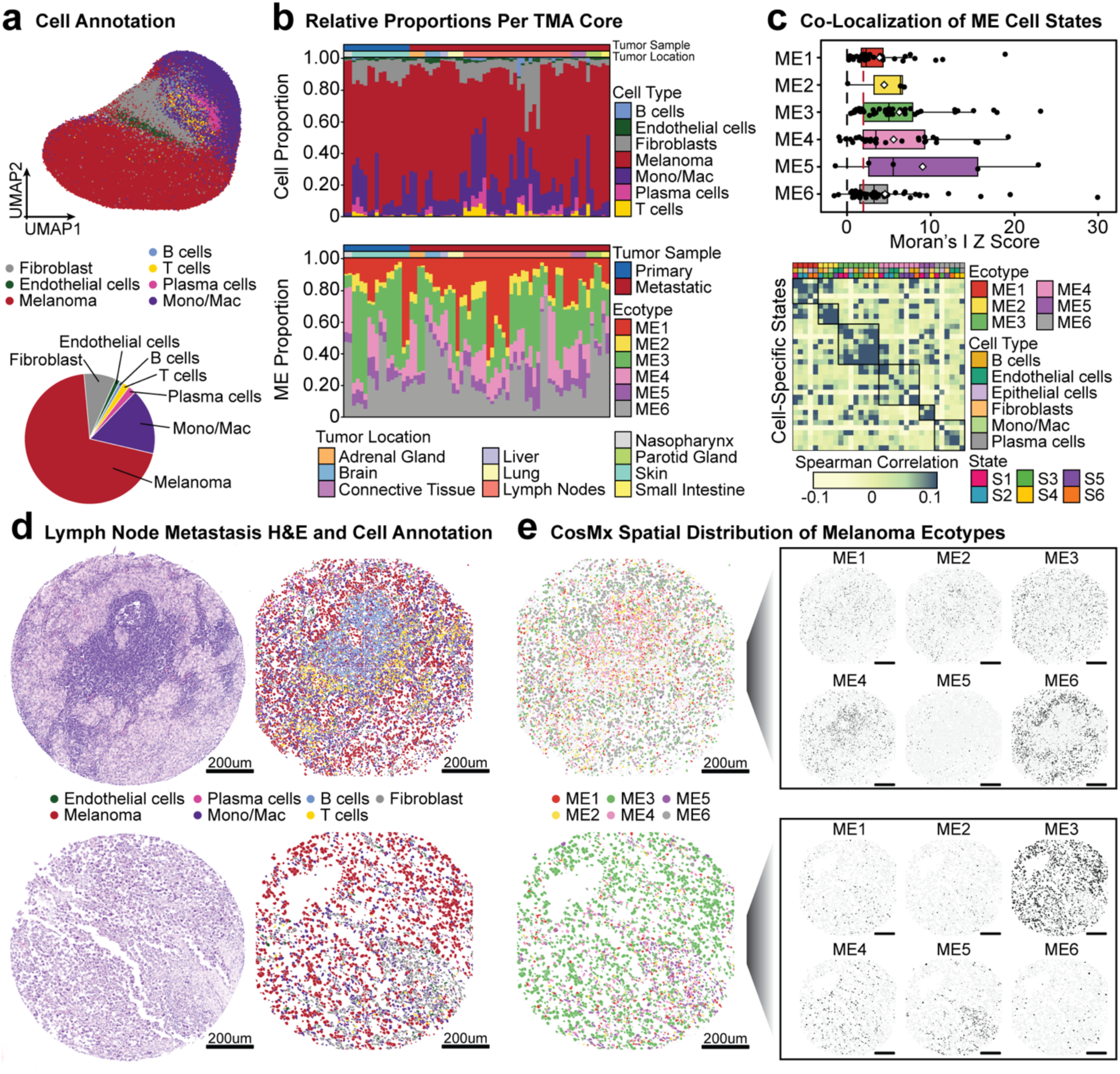
Spatial transcriptomic analysis of melanoma ecotypes. **(a)** Uniform manifold approximation and projection for dimension reduction (UMAP) of annotated cells from the melanoma TMA samples. **(b)** Relative cell proportion of the seven cell types across the primary and metastatic melanoma TMA samples. **(c)** Co-localization of MEs profiled by spatial transcriptomics with k=10 nearest neighbors and Moran’s I test. Significance determined by the red dashed line indicating a Z score=1.96 (p value=0.05). **(d)** Melanoma H&E samples (left) and respective spatial transcriptomic-analyzed melanoma TMA core (right). **(e)** Melanoma TMA cores visualized by organization of MEs in CosMx. Scale bar is 200 μm.

From our spatial autocorrelation analysis of the melanoma TMA samples, we found that in general, the MEs co-localized with each other more than cell types and states assigned to other MEs (**Figure 8c**). After confirming our annotations with H&E (Hematoxylin and Eosin) staining images and the annotated spatial transcriptomics (**Figure 8d**), we visually compared the spatial orientation of each assigned ecotype across the TMAs. Our co-localization results were consistent with our visual inspection of each TMA core, such that a single ME was typically dominant, followed by a secondary ME comprising a smaller population, and the remaining MEs were minimally represented (**Figure 8e**). Our findings from the spatial transcriptomics analysis further support our discovered MEs and their orientation in the tumor microenvironment.

### PhenoMapR links melanoma ecotype discovery to survival and immunotherapy outcomes

To further validate if our discovery MEs are indeed representative of their prognosis, we used an orthogonal computational modeling approach developed by our group termed PhenoMapR^32^. PhenoMapR allows us to identify genomic signatures correlated to clinical survival and immune checkpoint inhibitor (ICI) therapy response outcomes (**Figure 9a**). The PhenoMapR pipeline provides a semi-supervised computational approach that maps phenotypes associated with bulk gene expression data onto other genomics data (e.g., bulk RNA-seq, scRNA-seq, and spatial). This supports mapping of biological signatures from large-scale studies (such as ICI therapy response) onto our spatial transcriptomics data.

**Figure 9.**
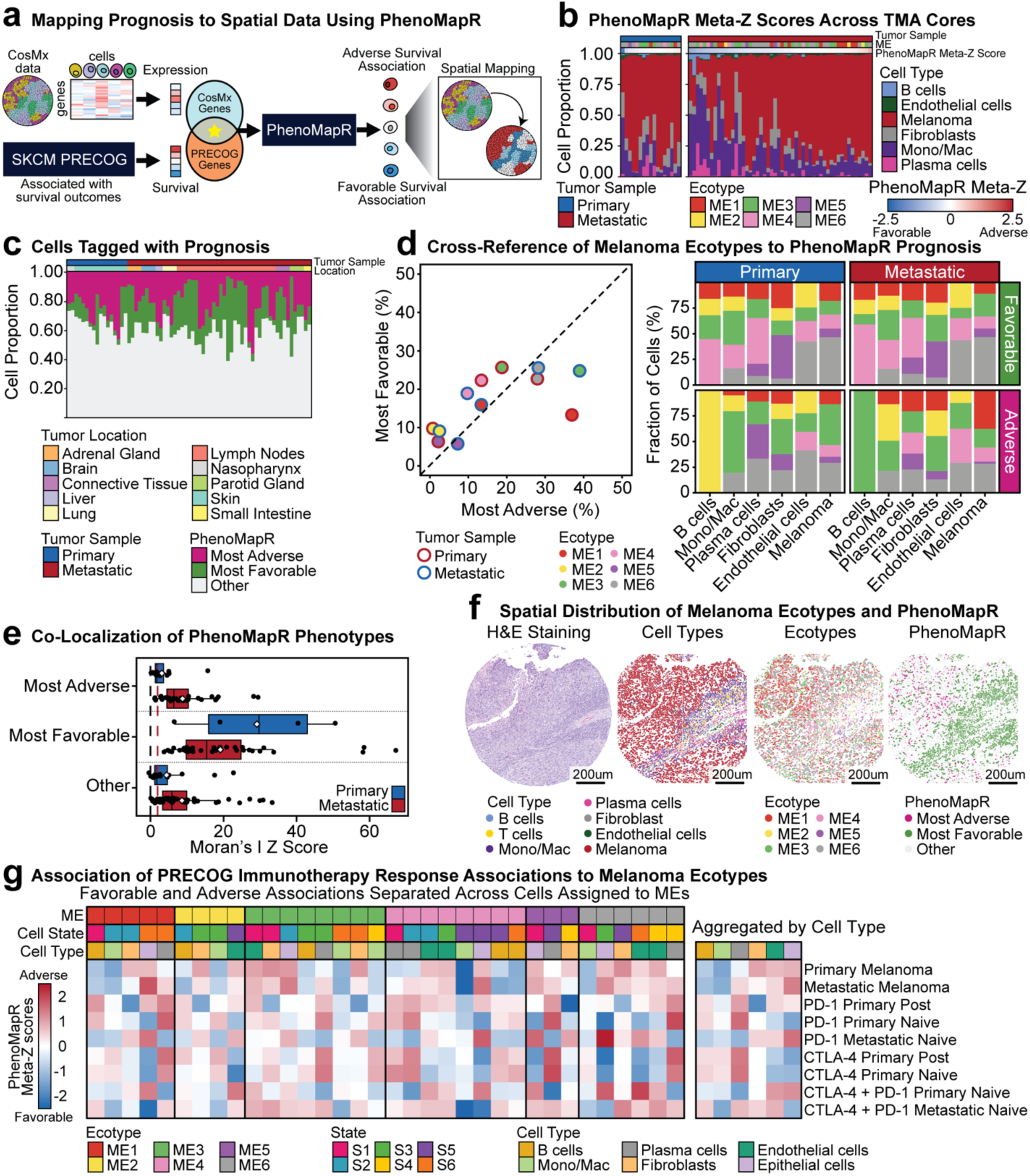
Application of PhenoMapR to identify phenotypes associated with favorable and adverse outcomes. **(a)** Computational overview of PhenoMapR to map phenotypes from bulk RNA-seq data from PRECOG onto spatial transcriptomics melanoma data. **(b)** Cell density across TMA cores ranked by their PhenoMapR meta-Z scores. Samples are separated by primary and metastatic diagnoses. **(c)** Distribution of cells assigned by PhenoMapR labels (most adverse, most favorable, and other) across all melanoma TMA cores. **(d)** Cross-reference of MEs and cell type distribution to phenotype assignment from PhenoMapR. **(e)** Co-localization of PhenoMapR phenotypes with k=10 nearest neighbors and Moran’s I test. Significance determined by the red dashed line indicating a Z score=1.96 (p value=0.05). **(f)** Representative melanoma H&E sample and respective spatial transcriptomics analyzed core annotated by cell types, MEs, and PhenoMapR phenotype labels. Scale bar represents 200 μm. **(g)** PhenoMapR meta-Z scores associated with melanoma PRECOG and ICI PRECOG clinical outcomes across the MEs. A positive meta-Z score is associated with adverse or non-responsive outcomes, whereas a negative meta-Z score is associated with favorable or response to ICI therapy.

We leveraged PhenoMapR to map survival outcomes from bulk RNA-seq primary and metastatic melanoma PRECOG meta-Z scores onto the spatial transcriptomic dataset^33,34^. After applying PhenoMapR to the melanoma CosMx dataset, we aimed to determine what cell types were contributing to the adverse and favorable survival outcomes. The cells with the top and bottom 20% association to the phenotypic survival outcomes (PhenoMapR meta-Z scores) were classified as most adverse and most favorable, respectively. Across the TMA cores, we found that the most adverse cores were represented by the melanoma, endothelial cells, and fibroblasts, while the most favorable cells were represented by the immune cells (**Figure 9b**). Comparing the cores for sample-specific bias, we found the cells tagged with PhenoMapR phenotypic survival outcomes to be uniformly distributed (**Figure 9c**).

We next cross-referenced our MEs and their respective cell types with the favorable and adverse phenotypic cell types determined by PhenoMapR. We found that primary and metastatic tumors with elevated ME4 ecotypes contained the highest proportion of favorable cells, whereas ME6 contained a greater number of cells with adverse associations (**Figure 9d**). Although ME2 was slightly associated with more favorable cells, its overall proportion was lower than that of ME4. The tumors assigned to ME1 and ecotypes contained with a greater number of cells with adverse associations in primary tumors, whereas those assigned to ME3 contained a greater number of cells in metastatic tumors. Both primary and metastatic tumors with higher ME5 abundance showed nearly equal proportions of adverse and favorable cells. These findings further demonstrate the favorable nature of ME4 and the heterogeneity of adverse associations among the other metastatic tumors.

We next compared the relative proportions of cells across MEs based on prognosis labeled by PhenoMapR (**Figure 9d**). Tumors abundant in ME1 contained more adverse melanoma and fibroblasts than other MEs. In contrast, ME2- and ME3-abundant tumors were characterized by more adverse immune cell types. We also observed a greater proportion of favorable cell type PhenoMapR labeling than adverse within tumors assigned to ME4. This finding further confirms our earlier TCGA SKCM finding that the abundant presence of immune cells in ME4 is associated with more favorable outcomes. Melanomas abundant in ME6 generally contained a similar proportion of adverse and favorable cell-type labeling. In summary, the metastasis-associated MEs (ME1-4 and ME6) and their differential clinical survival associations can be attributed to differences in their cellular composition and phenotypic associations with adverse outcomes. In contrast, the primary tumor-associated cellular community, ME5, indicates favorable outcomes if the plasma, fibroblast, and melanoma cells are labeled with favorable PhenoMapR scores.

To determine if these phenotypic labels spatially co-occurred together, we next performed spatial co-localization analysis for cells in each TMA, using their PhenoMapR label of most adverse, most favorable, and other. In general, we found that adverse cells were more aggregated in metastatic tumors, while favorable cells were more aggregated in primary tumors (**Figure 9e**). Among the TMAs, we were able to visualize the spatial orientation of the cells, along with MEs and the PhenoMapR phenotype classifications (**Figure 9f**).

With our discovery of six MEs having their own cell-type communities, enriched pathways, and survival outcomes, we asked if tumors assigned by their MEs responded differently to ICI therapy treatments. The ICI PRECOG database^24^ contains ICI outcomes on PD-1 (programmed cell death protein 1) inhibitors^35–40^, CTLA-4 (cytotoxic T-lymphocyte-associated protein 4) inhibitors^41–46^, and dual PD-1 + CTLA-4 inhibitors^35,47^. We compared these ICI treatments for SKCM in addition to untreated primary and metastatic melanoma PhenoMapR meta-Z scores to identify relative phenotypic shifts across MEs and treatments (**Figure 9g**). We included the primary and metastatic samples from PRECOG as a reference to individuals untreated with ICI therapy for SKCM. We found that favorable response signatures associated with CTLA-4 treatment of primary naïve and post-treated tumors assigned to ME1-2, ME4-5, and ME6. PD-1 treatment of primary naïve tumors showed favorable outcomes on tumors abundant with ME2 and ME4-6. ME3 exhibited the fewest favorable responses, with the strongest signal for CTLA-4 treatment of primary naïve tumors involving both epithelial and endothelial cells. The dual CTLA-4 + PD-1 treatment of metastatic naïve tumors displayed strong response signatures on tumors assigned to ME1, ME5, and ME6. Cell type importance in ICI response was most evident in epithelial melanoma cells, which exhibited a shift toward more favorable PhenoMapR meta-Z scores after mapping ICI treatment onto the spatial data. Together, these findings demonstrate concordant associations between the MEs and their constituent EcoTyper cell states with our PhenoMapR results, further supporting our identification of clinically relevant cellular communities associated with metastasis in melanoma.

## DISCUSSION

Our study aimed to identify the biological processes by which LN colonization by tumor cells promotes downstream distant metastasis in melanoma. Therefore, we hypothesized that integrating complementary pre-clinical and human tumor data would reveal relevant biology conserved across species.

We identified several cell cycle checkpoints, transcription, translation, and protein synthesis processes associated with melanoma linked with LN-colonization and metastasis. Disruption to the cell cycle checkpoints is a well-established sign of cancer^48^, and the G1/S transition mediated by cyclin-dependent kinase 4/6 is dysregulated in more than 96% of melanoma cell line cases^49^. Tumor cells that lack the G1/S checkpoints are known to rely on the G2/M checkpoint^50,51^, and our findings are consistent with this established dependency. Accordingly, therapeutic inhibition of this pathway^52,53^ or exploitation of this dependency through synthetic lethality^54,55^ may represent potential therapeutic strategies. Among the MEs, features of cell proliferation were enriched in the metastatic ME1 and depleted in primary tumor-associated ME5. In addition to cell cycle disruption, melanoma cells can induce DNA damage and stress signaling and develop drug resistance^56^. Relevant features of non-silent mutational rate were enriched in ME2 and depleted in ME5 and ME6. Alternatively, the number of altered chromosome arms and chromosomal instability was enriched in ME6. These MEs and their cellular communities provide further insight into the metastatic outcomes in humans.

Our PRECOG analysis helped validate the prognostic significance of the pathway-level and biomolecular differences identified between primary and metastatic tumors in our TransComp-R model. While the majority of pathways shared among the primary and metastatic conditions correlated with adverse or favorable outcomes, the TGFβ signaling pathway particularly stood out. The TGFβ signaling pathway plays a central role in tissue development, homeostasis, and repair^57^, and our findings suggest that genes pre-ranked by their PRECOG meta-Z scores were enriched for adverse outcomes under primary melanoma, but favorable in metastatic melanoma. Interestingly, others have reported similar conflicting properties of TGFβ acting as a tumor suppressor and stimulator of tumor metastasis^58–60^. For example, the activation of TGFβ signaling may promote tumorigenesis^61^, while inhibition may reduce metastatic abilities^62^. Others proposed TGFβ1 acts as a tumor promoter during the earlier stages of tumorigenesis^63^ and, during later stages, as a suppressor^64,65^ or stimulator^66,67^.

These discrepancies could be explained by our MEs, where the TGFβ feature is enriched in the neutral ME3, but depleted in primary-associated ME5 and the metastatic-associated ME6. In healthy cells, TGFβ can suppress tumor initiation and early-stage development by inhibiting cell cycle progression and suppressing growth factors^68^. The blockade of TGFβ signaling is favorable for immune cells (including T cells, B cells, NK cells, dendritic cells, and myeloid cells), but adverse when overactivated by cancer cells and fibroblasts in the tumor microenvironment^57,69,70^. This observation is likely since ME3 is characterized by elevated PMNs and fibroblasts, ME5 by mast and epithelial cells, and ME6 by endothelial cells. We noticed that ME3, enriched for TGFβ, is also enriched for epithelial-to-mesenchymal transition, whereas ME5, depleted of TGFβ signaling, showed depletion of epithelial-to-mesenchymal transition and proliferation^71^. Therefore, the tumor microenvironment balance between stromal and immune cells may contribute to the observed behaviors among our discovered MEs.

Our implementation of CIBERSORTx and EcoTyper allows us to establish melanoma-specific ecotypes. Notably, patients assigned to ME4 had the most favorable outcomes and would have been assigned to the favorable CE9 and CE10 from the pan-cancer carcinoma model^29^. The other MEs; however, did not overwhelmingly translate to a specific CE but rather distributed to more than one. Therefore, tumor samples that match similar cellular ME4 ecosystems may result in improved clinical outcomes compared to other MEs.

We also revealed cellular states unique to primary melanoma tumor samples in six cell types, including dendritic, endothelial, epithelial, mast, plasma cells, and PMNs. These cells also contained a panel of transcriptomic biomarkers predictive of LN metastasis outcomes that were expressed in primary but absent in metastatic cell states. Other groups have also found metastasis-associated features and behaviors of these cells. For instance, dendritic cells in the LN suppressed by melanoma cells were demonstrated to metastasize to distant sites^72^. Lymphatic endothelial cells have also been demonstrated to induce melanoma metastasis by driving chemotaxis of tumor cells^73,74^. Similar to other migratory cells, epithelial cells can undergo epithelial-to-mesenchymal transition and may enhance migratory capability and invasiveness^75–77^. Another notable immune cell type, mast cells, if residing in or around tumors, were associated with resistance to immunotherapy^78,79^. Melanoma with clusters of plasma cells resulted in significantly adverse survival outcomes compared to sparse or undetectable populations^80^. Finally, an elevated neutrophil-to-lymphocyte ratio indicated shortened outcomes in patients with melanoma^81,82^. Indeed, additional investigation of these cellular states across primary and metastatic conditions is warranted to determine clinical relevance.

Our spatial transcriptomics analysis confirmed that our defined cell types and states within each ME indeed co-occur within the tumor microenvironment. We further validated these findings using an orthogonal computational framework termed PhenoMapR, which mapped phenotypes associated with favorable and adverse clinical outcomes from bulk expression data onto our spatial transcriptomics data. These findings together reinforce the biological relevance of the discovered MEs, and they discover clinically informative tumor ecosystems in melanoma.

This study contains limitations. For example, the number of genes analyzed was inevitably limited to cross-species homolog pairs. The TransComp-R pipeline does not account for all genes across different species; therefore, some genes that may be relevant in the human or mouse cohorts may be excluded by the requirement for one-to-one gene match during the matrix multiplication step. Although LN colonization is not unique to melanoma and likely extrapolates a broader behavior of colonization across all solid tumors^1,2,83^, our results are derived from melanoma-specific studies and should be generalized with caution.

Our systems biology approach allowed us to identify significantly enriched biological pathways that link together LN colonization and metastatic outcomes in melanoma through translational modeling of mouse to human. We further interpreted our findings and identified biomolecular and pathway-level associations with survival outcomes and identified distinct MEs defined by their cell states and prognosis. We then performed spatial autocorrelation analysis and confirmed these MEs co-occur together in melanoma TMAs. Indeed, these findings reveal a potential connection between the process by which tumors colonize the LNs and metastasize throughout the body.

## MATERIALS AND METHODS

### Bulk RNA sequenced data selection

We accessed bulk RNA-seq tumor samples from GEO that contained gene expression. We used search keywords including combinations of “melanoma,” “RNA-seq,” “primary,” “metastasis,” and “gene expression.” We also aimed to prioritize datasets with balanced samples and recorded demographic or clinical variables.

### Pre-processing and normalization

We accessed human transcriptomics data deposited in the GEO repository by using the GEO2R pipeline in R. (GEOquery v2.76.0, Biobase v2.68.0, and limma v3.64.3)^84–86^. We log_2_-transformed the data and Z scored the data per gene. In preparation for the TransComp-R workflow, we matched shared human homolog pairs between the mouse and human data using orthogene (v1.14.0)^87^. The genes that did not contain an exact one-to-one match were omitted from the computational model and from further analysis.

### TransComp-R cross-species modeling

To synthesize the mouse and human data together, we first performed PCA on the mouse dataset. We filtered for genes that contributed to a maximum cumulative explained variance of 80% to prevent overfitting of the data. We then performed TransComp-R by multiplying the matrix ***X*** with ***s*** rows of human samples and ***n*** columns of genes by the ***Q*** matrix containing ***n*** rows of genes and ***p*** columns of mouse LN PCs. The resulting product becomes a projected matrix ***P*** with ***s*** rows of human subjects and ***p*** columns of mouse LN PCs (**Eq. 1**).

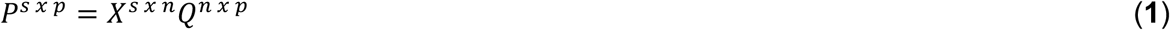

### Principal component feature selection

We leveraged LASSO to select the mouse LN PCs predictive of human metastatic outcomes. To address potential confounders, we incorporated variables of sex, age, and interaction terms. Our LASSO model was established using 100 random rounds of 10-fold cross-validation regressing against human outcomes. The regression model can be described with **Eq. 2**:

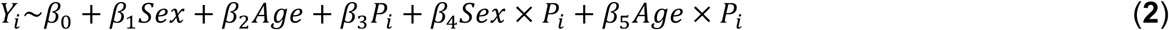

such that ***Y*** is the human outcomes classified by metastasis or primary, ***Sex*** and ***Age*** representing the human demographic information, and ***P_i_*** representing the projected human matrix for the ***i^th^*** PC-human projection. A model p value less than 0.05 was established for significance of a particular mouse LN PC.

### Percent variance explained across species

We calculated the variance explained in the human data by each mouse PC to determine the translational potential across mouse to the human space. We can compute the variance explained in human data using the human melanoma tumor data matrix ***X*,** matrix ***Q*** containing columns of mouse LN PCs, and each PC, ***q_i,_*** from the ***Q*** matrix. The ***T*** represents the transposed matrix to calculate the variance explained in human by mouse (**Eq. 3**).

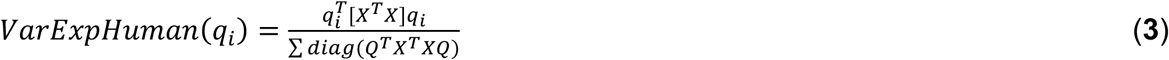

### Gene set enrichment analysis

We pre-ranked genes by their scores defined by each PC of interest and identified enriched biological pathways with GSEA^88^ on R (fgsea v1.34.2 and clusterProfiler v4.16.0)^89,90^. We referenced the KEGG and Hallmark curations for a holistic background on the biological pathways from msigdbr (v25.1.1)^91,92^. Our parameters for the analysis included a minimum gene size of 5, a maximum gene size of 500, and an epsilon coefficient of 0. We used the default 1,000 permutations and defined a pathway to be significant with a Benjamini-Hochberg FDR value less than 0.05.

### Gene association to prognosis

We used the PRECOG platform (precog.stanford.edu)^23,24^ to match genes encoded on the mouse LN PC of interest with meta-Z scores indicative of cancer prognosis. We compared meta-Z scores between the metastatic and primary outcomes of melanoma to the PC1 loadings and their respective PC scores. We then pre-ranked genes encoded on the selected mouse PCs of interest and ran GSEA with the same input parameters of a minimum gene size of 5, maximum gene size of 500, epsilon coefficient of 0, and the default 1,000 permutations. All pathways were determined to be significant if their Benjamini-Hochberg adjusted FDR values were less than 0.05.

### Bulk RNA-sequenced cell type deconvolution

We deconvolved bulk RNA-seq data using the CIBERSORTx platform (cibersortx.stanford.edu)^28^, which allows for the estimation of specific cell types using bulk RNA-seq data. With CIBERSORTx, we used two signature matrices to more accurately quantify the cellular composition in the tumor microenvironment. We first used the TR4 signature matrix to first estimate endothelial (CD31+), epithelial (EPCAM+), fibroblast (CD10+), and pan-immune (CD45+) cell populations derived from resected tumor samples from patients with non-small cell lung cancer^93,94^.

To characterize the distinct immune cells, we used the LM22 signature matrix, which contains an established number of genes that defines the cell expression profile of 22 different immune cells. We used the LM22 signature matrix and merged related subpopulations into 10 distinct immune cells for interpretability, robustness, and statistical power^28,94^. These 10 immune cells included B cells, CD4+ T cells, CD8+ T cells, dendritic cells, eosinophils, mast cells, monocytes and macrophages, NK cells, plasma cells, and PMN cells. With CIBERSORTx, we ran each deconvolution model with 1,000 permutations and disabled quantile normalization.

### In silico cell type fraction estimation

After estimating the four broad cell types with the TR4 signature matrix, we further refined the pan-immune fraction using the LM22 signature matrix. We concomitantly integrated the 10 immune cells from the LM22 signature matrix by proportionally allocating the pan-immune fraction to each sample and its cellular fractions, such that all cellular components totaled to a summed relative ratio of 1 per sample. This process resulted in the estimated cellular proportions of 13 cells, including endothelial, epithelial, fibroblasts, B cells, CD4+ T cells, CD8+ T cells, dendritic cells, eosinophils, mast cells, monocytes and macrophages, NK cells, plasma cells, and PMN cells. This approach was also repeated for the EcoTyper model with the exclusion of eosinophils to allow for equal comparison of the same cellular components for model evaluation to previously established EcoTyper models in carcinoma^29^.

### High-resolution expression imputation

To calculate specific gene expression in the cell types of interest, we used the estimated cell type abundances from CIBERSORTx to purify the cell-specific gene expression. We disabled quantile normalization for the high-resolution analysis and maintained all other default settings. The combined TR4 and LM22 signature matrix was used for high-resolution expression imputation.

### Tumor ecosystems and cell state discovery

We leveraged discovery mode in the EcoTyper pipeline^29^ to define the MEs. To cross-reference our findings to a previously established carcinoma EcoTyper discovery model, we deconvolved the bulk SKCM RNA-seq data from the TCGA to the same 12 major cell types presented in the established carcinoma EcoTyper model^29^, which included B, CD4+ T, CD8+ T, dendritic, endothelial, epithelial, fibroblast, mast, monocytes and macrophages, NK, PMN, and plasma cells.

We extracted cell type genes by matching genes with a non-zero variance present in our calculated high-resolution and carcinoma cell profiles in the bulk RNA-seq data matrix and performed non-negative matrix factorization. We defined the number of threads to be 12, and the number of non-negative matrix factorization restarts to 5. We used a cophenetic coefficient cutoff of 0.95, a maximum number of states per cell type of 20, and a minimum number of states in an ecotype of 3. After we selected the number of cell states and performed quality control filtering, we generated the discovery for MEs.

### Co-occurrence calculation across melanoma ecotype networks

We constructed a visualized network of the cell states for each ME in the igraph (v2.1.4)^95^ package and prepared the network in Cytoscape (v3.10.4)^96^. We proportionally weighted the edges between each cell state node determined by its respective Jaccard index between each cell state. This network presents the degree of overlap between each cell state across the tumor samples in the discovery cohort.

### Survival analysis

We performed survival analysis in R using the package survminer (v0.5.1) on patients classified by their ME. A log-rank test was performed to compare significance across the groups.

### Carcinoma ecotype recovery

We aimed to compare the MEs to the previously validated CEs with the goal of identifying sample overlaps across the two ecotype models. We generated the recovery ecotype model with default parameters and evaluated the different MEs with an alluvial diagram tracing the sample identifiers across both carcinoma and melanoma ecotypes.

### Analysis of biological features across melanoma ecotypes

To summarize the MEs and their associated biological features, we used curated data from the Hallmark MSigDB and Immunogenomics features from Thorsson et al.^31^ to link genomic characteristics to biological associations in tumor samples. We calculated enrichment of the biological features in the bulk TCGA SKCM tumors by their most abundant ME. We determined the enrichment or depletion using a two-sided Wilcoxon test by comparing each ME relative to other MEs in the TCGA SKCM cohort. We adjusted the p values for multiple hypothesis testing using the Benjamini-Hochberg method, and features with an FDR<0.05 were considered significantly enriched or depleted in the ME.

### Spatial transcriptomics tissue microarray preparation

Melanoma tissue samples were acquired from the Stanford Cancer Institute Tissue Bank. All sample collections in this study were collected with informed consent for research. All research protocols were approved by the Stanford University Institutional Review Board and performed in strict accordance with the Declaration of Helsinki.

The TMA was prepared and stained following the guidelines in the CosMx SMI Manual Slide Preparation for RNA Assays manual (MAN-10184-06). The FFPE tissue block was sectioned at 5 μm and mounted on LEICA Bond Plus slides. The default experimental conditions were used for target retrieval time, digestion buffer concentration, and digestion time. Fiducials were applied at a concentration of 0.0005% per manufacturer’s protocol manual for skin cancer tissue. Tissue sections were hybridized overnight with the CosMx Human 6k Discovery Panel and stained the next day with CosMx DAPI nuclear stain, CosMx Hs CD298/B2M, and CosMx Hs PanCK/CD45 markers. Pre-bleaching profile configuration C was selected during imaging on the CosMx machine per manufacturer’s recommendation.

### Cell segmentation

Cell segmentation was performed using the AtoMx Spatial Informatics Platform by Bruker. We segmented the cells using the manual’s recommended configuration A (non-neuro human tissue) with default nucleus label model bsbNuc and cytoplasm label model bsbCyto. The parameters include nuclear diameter (6.64 μm), cell diameter (7.71 μm), minimum cell size (3.51 μm), and cell dilation (0.4 μm). All cores were then visually inspected to confirm accurate segmentation.

### Spatial transcriptomics data processing

We performed all spatial transcriptomics analysis in R with Seurat packages^97^. We removed cells exceeding the 99^th^ percentile of nCounts and nFeatures and removed cells with a nCount less than 50 and nFeature of 150. We removed FOVs with fewer than 100 cells as recommended for 6k-plex CosMx kits by Bruker Corporation. We log-normalized the samples with a scale factor of 10,000. We found variable features with nFeatures of 2,000 genes and 30 PCs. We used harmony with a theta of 0.05 to adjust for unwanted variation across different donors. We found clusters using a resolution of 1.2. Next, we performed a UMAP with 12 dimensions, harmony reduction, Euclidian distance, and 15 nearest neighbors.

### Cell type annotation

We first used Insitutype (v2.0)^98^ to annotate the Seurat clusters. We used supervised learning with Insitutype using the TR4 gene expression profile to determine cells that were epithelial, endothelial, fibroblast, and pan immune cells. We then used Insitutype with semi-supervised learning with the LM22 signature matrix with the option of 12 new clusters to annotate the immune cells. Using these two annotation files, we superimposed the TR4 annotations onto the LM22 annotations such that the non-immune cells were annotated by TR4, and immune cells were annotated by LM22.

We next used FindAllMarkers on the cell type clusters identified by Insitutype with genes detected in at least 25% of the cells. We used CyteTypeR (v0.4.2)^99^ using the top 10 gene markers of each cluster to further interpret the cell annotations. After combining shared cell types into new clusters, we iteratively subclustered each cell cluster through the Seurat pre-processing steps and reassigned subgroups to matching cell types. This process was repeatedly performed until the cell annotations for each cell type clusters were labeled.

We determined cell type annotations with the following gene expression markers: Melanoma (*MLANA*, *PMEL*, *SOX10*), T cells (*CD3D*, *CD3E*, *CD3G*, *ITK*, *IL7R*), B cells (*MS4A1*, *CD79A*, *CD19*), monocytes and macrophages (*C1QA*, *C1QB*, *HLA-DRA*, *CD74*, *CD68*, *CD163*), plasma cells (*IGHG1/2*, *IGLC1/2*, *JCHAIN*, *IGHA1*), fibroblast (*COL1A1*, *COL1A2*, *DCN*) and endothelial cells (*PECAM1*, *VWF*).

### Spatial autocorrelation analysis

We performed a Moran’s I test to calculate the aggregation patterns of cell types and states assigned to MEs^100^. We used the cells assigned to the most abundant ME from the spatial transcriptomics recovery from our discovery model. These assigned MEs were used to generate spatial transcriptomic visualization and co-localization analysis with a Spearman correlation analysis and Moran’s I test. The Moran’s I test is calculated using **Eq. 4**.

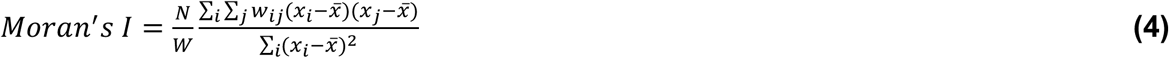

Here, ***N*** represents the total number of cells, ***x*** is the score for each ME, ***x***A is the mean of ***x***, and ***w_ij_*** are the spatial weights. ***W*** is the sum of the spatial weights.

To account for TMA cores that may have a scarce number of cells, we performed a low-grade filter such that each TMA must contain at least 300 cells per core, 300 cells per ecotype, and an assigned ecotype consisting of at least 10% of the total number of cells. TMA cores containing values less than these input values for a given ME calculation were not included for the Moran’s I test.

### Determination of adverse and favorable clinical outcomes of melanoma samples

We used the PhenoMapR (v0.1.0) to map melanoma-relevant phenotypes from bulk expression onto spatial transcriptomics data^32^. We used the pre-loaded database of prognostic meta-Z scores from PRECOG and ICI PRECOG specific to SKCM. The PhenoMapR scores for each pre-loaded database were computed using the processed melanoma TMA CosMx Seurat object as the input file. The calculated PhenoMapR score from each data source was scaled to the same minimum and maximum value while also preserving the original zero value. We then performed Z score normalization across the PhenoMapR PRECOG and ICI PRECOG variables across all MEs to compute a relative meta-Z score indicating adverse and favorable outcomes in survival and therapeutic responses.

## ACKNOWLEDGEMENTS

This work is supported by the National Cancer Institute Cancer Systems Biology Consortium under grant number U54CA274511 and National Cancer Institute R01CA276828 (AJG). BKB is supported by the Propel Postdoctoral Fellowship.

## AUTHOR CONTRIBUTIONS

**BKB:** Conceptualization, data curation, formal analysis, investigation, methodology, visualization, writing-original draft, writing-review & editing. **RA:** Data curation, writing-review & editing. **SZ:** Data curation, writing-review & editing. **II:** Data curation, writing-review & editing. **BAB:** Data curation, writing-review & editing. **BEH:** Data curation, resources, writing-review & editing. **AJG:** Conceptualization, funding acquisition, methodology, project administration, resources, writing-review & editing.

## DATA AND CODE AVAILABILITY

We accessed bulk RNA-sequenced data from Gene Expression Omnibus with accession numbers GSE117529, GSE46517, and GSE65904. Additional bulk RNA-sequenced SKCM data was accessed from The Cancer Genome Atlas. The melanoma CosMx spatial transcriptomics data will be made publicly available on Gene Expression Omnibus upon publication. All code for the analysis is deposited and made publicly available on GitHub: https://github.com/Gentles-lab/MouseToHumanMelanoma. The source code for the EcoTyper model was accessed on GitHub: https://github.com/digitalcytometry/ecotyper. PhenoMapR was accessed on GitHub: https://github.com/brooksbenard/PhenoMapR.

## SUPPLEMENTARY FIGURES

**Supplementary Figure S1.**
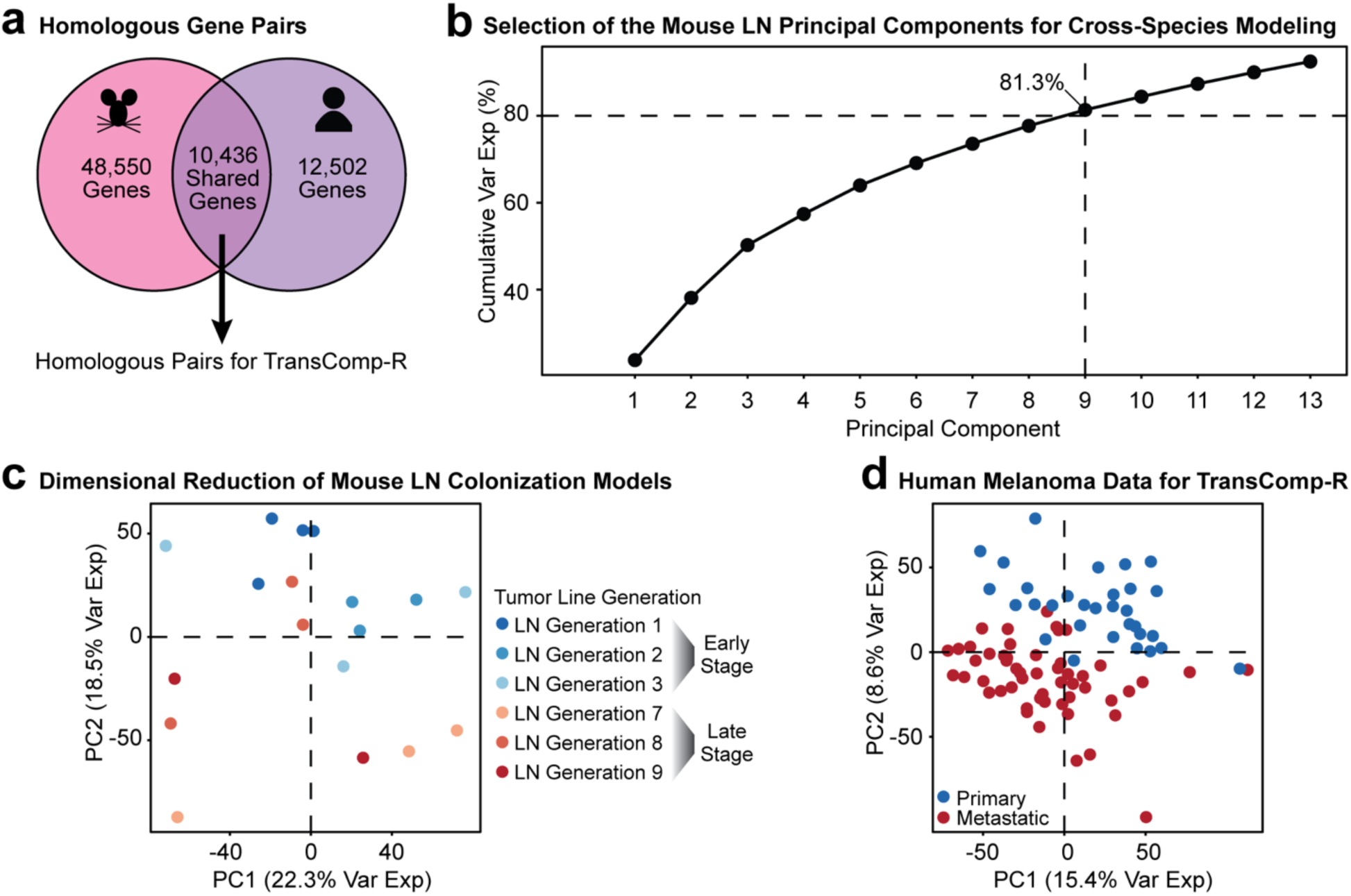
Pre-processing of mouse and human bulk RNA-seq data. **(a)** Matched homologs between mouse and human data. **(b)** Nine PCs explain 80% of the cumulative variance in mouse. **(c)** PCA of the mouse LN colonization samples separated by the early- and late-stage generations (percent variance explained in mouse). **(d)** PCA of the human melanoma data used for the TransComp-R pipeline (percent variance explained in human).

**Supplementary Figure S2.**
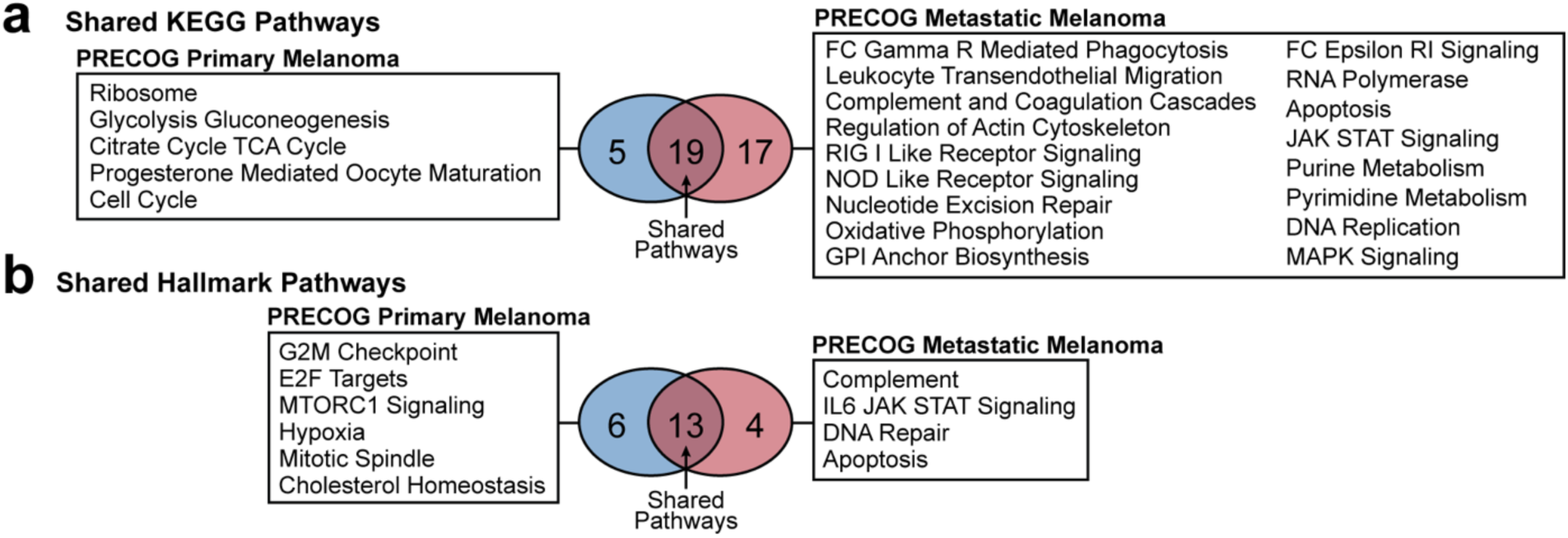
Shared enriched pathway across PRECOG primary and metastatic melanoma. Comparing the number of overlapping pathways across the primary and metastatic melanoma samples pre-ranked by PRECOG meta-Z scores for both **(a)** KEGG and **(b)** hallmark curations.

**Supplementary Figure S3.**
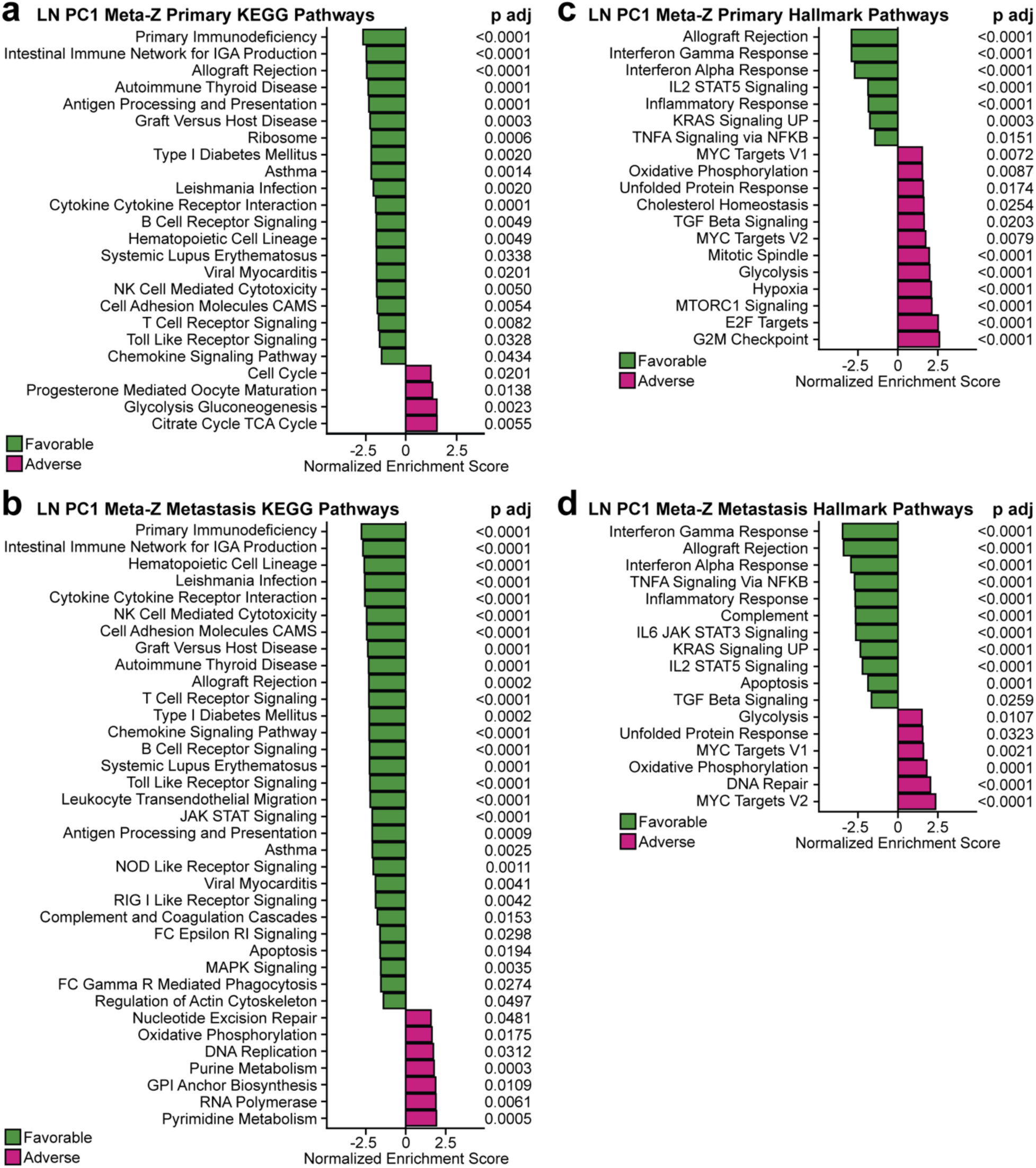
Pathway enrichment analysis of the mouse LN PC1 re-ranked by PRECOG meta-Z scores. Enriched KEGG pathways on LN PC1 for **(a)** primary melanoma and **(b)** metastatic melanoma. Enriched hallmark pathways on LN PC1 for **(c)** primary melanoma and **(d)** metastatic melanoma. The p values are adjusted by the Benjamini-Hochberg method.

**Supplementary Figure S4.**
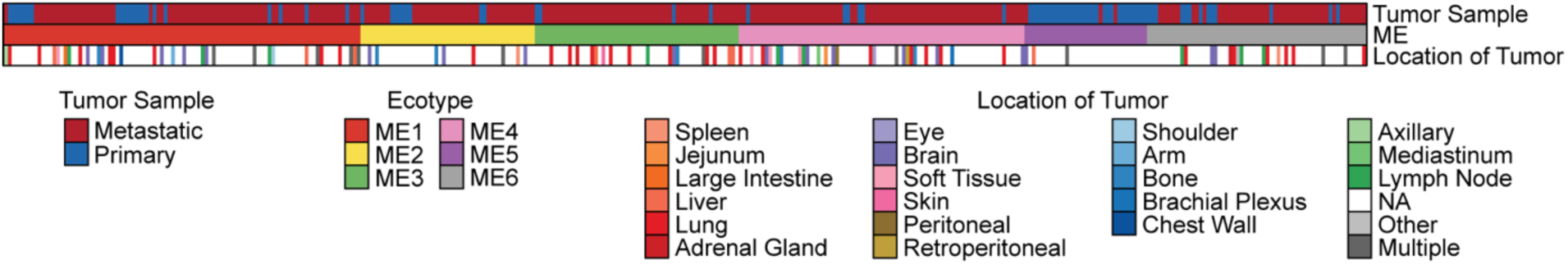
Breakdown of tumor location with respect to each melanoma ecotype from the discovery model. The tumor sample, melanoma ecotype, and location of the resected tumor are shown from the TCGA SKCM cohort.

**Supplementary Figure S5.**
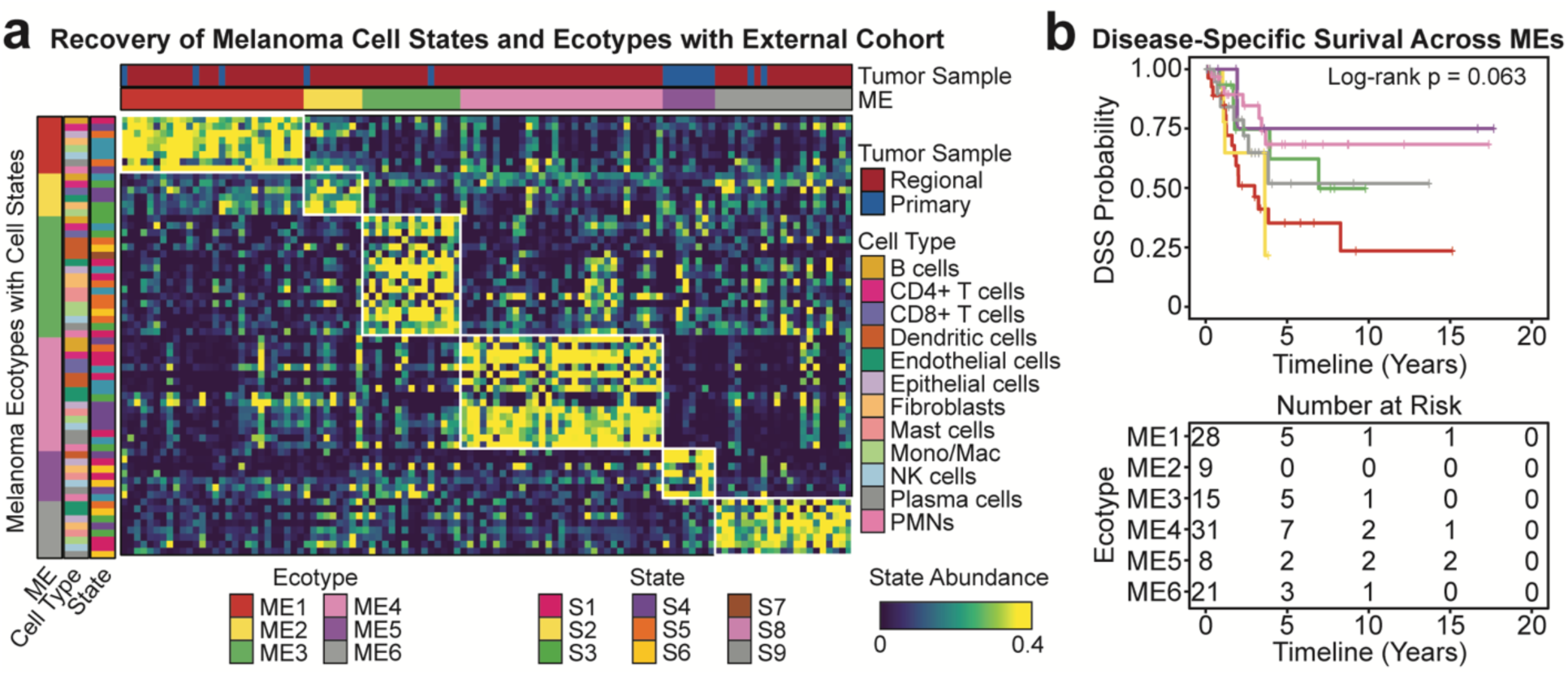
External validation of the melanoma ecotypes in GSE65904. **(a)** Recovered MEs and cell states. **(b)** Disease-specific survival of patients aggregated by MEs.

**Supplementary Figure S6.**
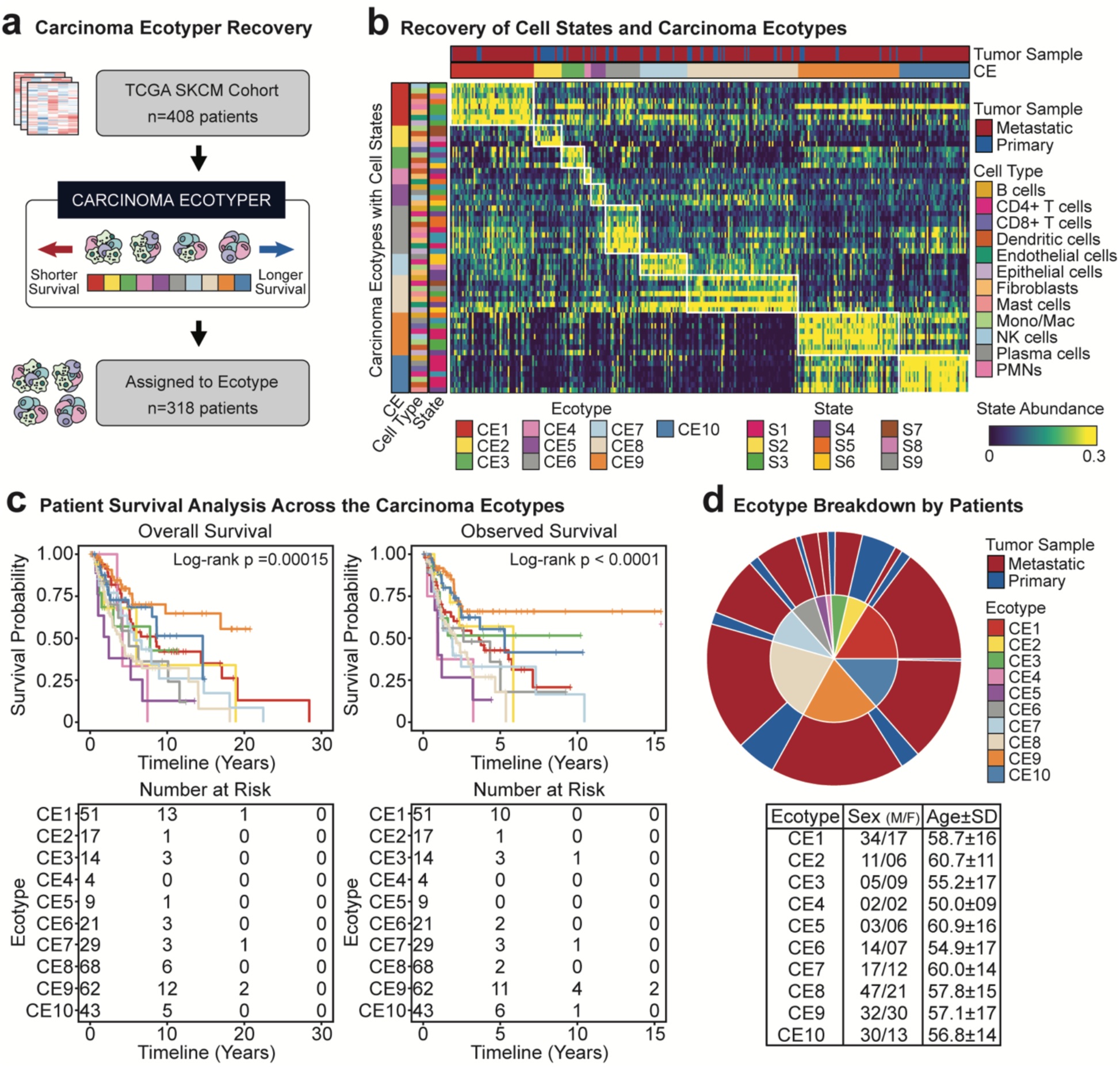
Recovery of carcinoma ecotypes. **(a)** Recovered carcinoma ecotypes using the TCGA SKCM dataset. **(b)** Distribution of carcinoma ecotypes across the tumor samples (n=318 assigned to a CE). **(c)** Patient survival outcomes defined by overall survival and observed survival. **(d)** Ecotype breakdown by patient demographics. Age is displayed with mean age and the standard deviation for each carcinoma ecotype.

**Supplementary Figure S7.**
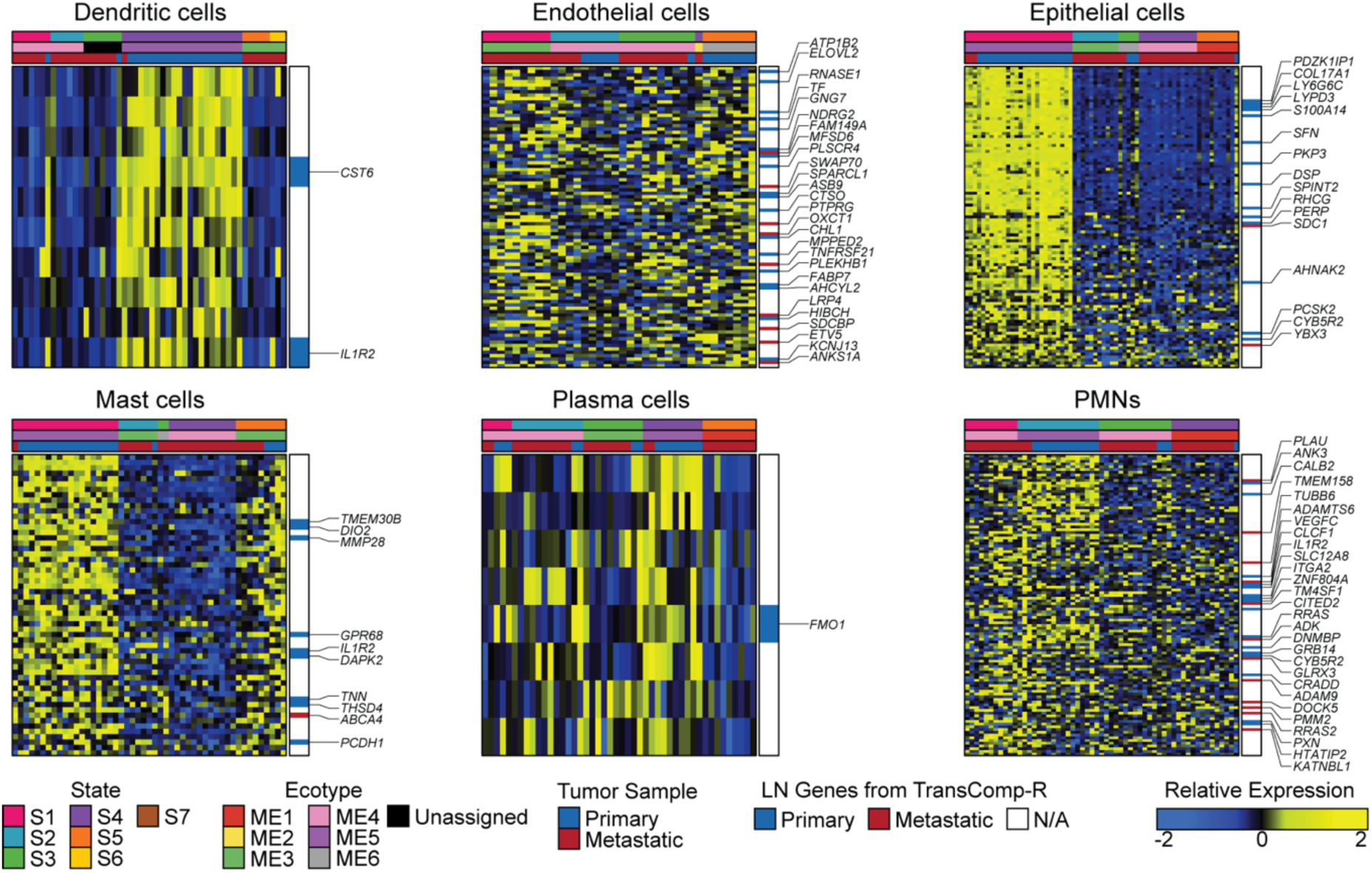
Gene subset of the recovered cell states unique to a primary tumor-specific state. Cell-state expression annotated by genes predictive of LN metastasis by TransComp-R.

**Supplementary Figure S8.**
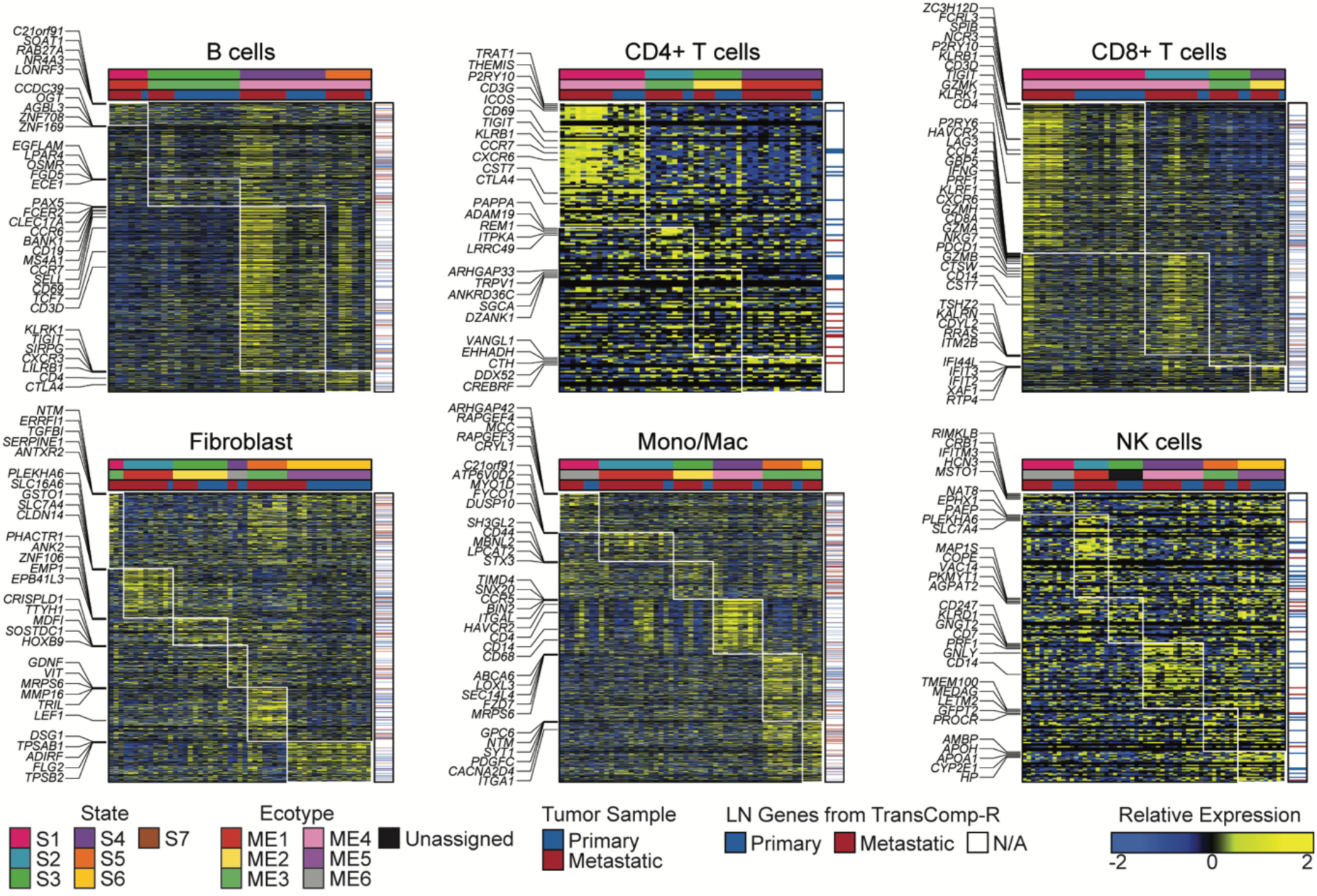
Additional cell-specific gene expressions on other differentially abundant cells from recovered cells in GSE46517. Each cell is annotated by its cellular state, tumor sample type, and genes that have predictive ability of LN metastasis as identified by TransComp-R.

## REFERENCES

1. Ji, H. et al. Lymph node metastasis in cancer progression: molecular mechanisms, clinical significance and therapeutic interventions. Signal Transduct. Target. Ther. 8, 367 (2023).

2. Pereira, E. R., Jones, D., Jung, K. & Padera, T. P. The lymph node microenvironment and its role in the progression of metastatic cancer. Semin. Cell Dev. Biol. 38, 98–105 (2015).

3. Alitalo, K. The lymphatic vasculature in disease. Nat. Med. 17, 1371–1380 (2011).

4. Brown, M. et al. Lymph node blood vessels provide exit routes for metastatic tumor cell dissemination in mice. Science 359, 1408–1411 (2018).

5. Haist, M. et al. Lymph node colonization induces tissue remodeling via immunosuppressive fibroblast-myeloid cell niches supporting metastatic tolerance. Cancer Cell 0, (2026).

6. Reticker-Flynn, N. E. et al. Lymph node colonization induces tumor-immune tolerance to promote distant metastasis. Cell 185, 1924–1942.e23 (2022).

7. Riedel, A., Shorthouse, D., Haas, L., Hall, B. A. & Shields, J. Tumor-induced stromal reprogramming drives lymph node transformation. Nat. Immunol. 17, 1118–1127 (2016).

8. Farzad, Z. et al. Lymphocyte Subset Alterations in Nodes Regional to Human Melanoma1. Cancer Res. 50, 3585–3588 (1990).

9. Ruddell, A., Harrell, M. I., Furuya, M., Kirschbaum, S. B. & Iritani, B. M. B Lymphocytes Promote Lymphogenous Metastasis of Lymphoma and Melanoma. Neoplasia 13, 748–757 (2011).

10. Lee, S. Y. et al. Changes in specialized blood vessels in lymph nodes and their role in cancer metastasis. J. Transl. Med. 10, 206 (2012).

11. Yang, W. et al. Targeting high endothelial venules: potential strategies for cancer treatment. Ann. Med. 57, 2597587 (2025).

12. Mestas, J. & Hughes, C. C. W. Of Mice and Not Men: Differences between Mouse and Human Immunology. J. Immunol. 172, 2731–2738 (2004).

13. Rangarajan, A. & Weinberg, R. A. Comparative biology of mouse versus human cells: modelling human cancer in mice. Nat. Rev. Cancer 3, 952–959 (2003).

14. Rivera, J. & Tessarollo, L. Genetic Background and the Dilemma of Translating Mouse Studies to Humans. Immunity 28, 1–4 (2008).

15. Mak, I. W., Evaniew, N. & Ghert, M. Lost in translation: animal models and clinical trials in cancer treatment. Am. J. Transl. Res. 6, 114–118 (2014).

16. Heng, H. H. et al. Heterogeneity Mediated System Complexity: The Ultimate Challenge for Studying Common and Complex Diseases. in The Value of Systems and Complexity Sciences for Healthcare (ed. Sturmberg, J. P.) 107–120 (Springer International Publishing, Cham, 2016). doi:10.1007/978-3-319-26221-5_9.

17. Brubaker, D. K. et al. An interspecies translation model implicates integrin signaling in infliximab-resistant inflammatory bowel disease. Sci. Signal. 13, eaay3258 (2020).

18. Ball, B. K., Park, J. H., Bergendorf, A. M., Proctor, E. A. & Brubaker, D. K. Translational disease modeling of peripheral blood identifies type 2 diabetes biomarkers predictive of Alzheimer’s disease. Npj Syst. Biol. Appl. 11, 1–16 (2025).

19. Frost, M. R. et al. Computational translation of mouse models of osteoarthritis predicts human disease. Osteoarthritis Cartilage 0, (2025).

20. Ball, B. K. et al. Integrated cross-species translation and biophysical multi-scale modeling links molecular signatures and locomotory phenotypes in spaceflight-induced sarcopenia. Npj Microgravity 10.1038/s41526-025-00557-x (2026) doi:10.1038/s41526-025-00557-x.

21. Kabbarah, O. et al. Integrative Genome Comparison of Primary and Metastatic Melanomas. PLOS ONE 5, e10770 (2010).

22. Tibshirani, R. Regression Shrinkage and Selection Via the Lasso. J. R. Stat. Soc. Ser. B Methodol. 58, 267–288 (1996).

23. Gentles, A. J. et al. The prognostic landscape of genes and infiltrating immune cells across human cancers. Nat. Med. 21, 938–945 (2015).

24. Benard, B. A., Lalgudi, C. K., Ilerten, I., Wang, R. H. & Gentles, A. J. PRECOG update: an augmented resource of clinical outcome associations with gene expression for adult, pediatric, and immunotherapy cohorts. Nucleic Acids Res. 54, D1579–D1589 (2026).

25. Pei, H. et al. FKBP51 Affects Cancer Cell Response to Chemotherapy by Negatively Regulating Akt. Cancer Cell 16, 259–266 (2009).

26. Li, L., Lou, Z. & Wang, L. The role of FKBP5 in cancer aetiology and chemoresistance. Br. J. Cancer 104, 19–23 (2011).

27. Marrone, L. et al. Exploring the potential of selective FKBP51 inhibitors on melanoma: an investigation of their in vitro and in vivo effects. Cell Death Discov. 11, 138 (2025).

28. Newman, A. M. et al. Robust enumeration of cell subsets from tissue expression profiles. Nat. Methods 12, 453–457 (2015).

29. Luca, B. A. et al. Atlas of clinically distinct cell states and ecosystems across human solid tumors. Cell 184, 5482–5496.e28 (2021).

30. Xiong, J., Bing, Z. & Guo, S. Observed Survival Interval: A Supplement to TCGA Pan-Cancer Clinical Data Resource. Cancers 11, (2019).

31. Thorsson, V. et al. The Immune Landscape of Cancer. Immunity 48, 812–830.e14 (2018).

32. Benard, B. A., Lalgudi, C. K., Azizi, A. & Gentles, A. J. PhenoMapR: scalable mapping of sample phenotypes to single-cell, spatial, and bulk transcriptomics data. 2026.09.08.749933 Preprint at 10.64898/2026.09.08.749933 (2026).

33. Cirenajwis, H. et al. Molecular stratification of metastatic melanoma using gene expression profiling: Prediction of survival outcome and benefit from molecular targeted therapy. Oncotarget 6, 12297–12309 (2015).

34. Budden, T. et al. Repair of UVB-induced DNA damage is reduced in melanoma due to low XPC and global genome repair. Oncotarget 7, 60940–60953 (2016).

35. Gide, T. N. et al. Distinct Immune Cell Populations Define Response to Anti-PD-1 Monotherapy and Anti-PD-1/Anti-CTLA-4 Combined Therapy. Cancer Cell 35, 238–255.e6 (2019).

36. DeVito, N. C. et al. Pharmacological Wnt ligand inhibition overcomes key tumor-mediated resistance pathways to anti-PD-1 immunotherapy. Cell Rep. 35, 109071 (2021).

37. Riaz, N. et al. Tumor and Microenvironment Evolution during Immunotherapy with Nivolumab. Cell 171, 934–949.e16 (2017).

38. Cui, C. et al. Ratio of the interferon-γ signature to the immunosuppression signature predicts anti-PD-1 therapy response in melanoma. NPJ Genomic Med. 6, 7 (2021).

39. Prat, A. et al. Immune-Related Gene Expression Profiling After PD-1 Blockade in Non-Small Cell Lung Carcinoma, Head and Neck Squamous Cell Carcinoma, and Melanoma. Cancer Res. 77, 3540–3550 (2017).

40. Liu, D. et al. Integrative molecular and clinical modeling of clinical outcomes to PD1 blockade in patients with metastatic melanoma. Nat. Med. 25, 1916–1927 (2019).

41. Wolchok, J. D. et al. Overall Survival with Combined Nivolumab and Ipilimumab in Advanced Melanoma. N. Engl. J. Med. 377, 1345–1356 (2017).

42. Van Allen, E. M. et al. Genomic correlates of response to CTLA-4 blockade in metastatic melanoma. Science 350, 207–211 (2015).

43. Chen, P.-L. et al. Analysis of Immune Signatures in Longitudinal Tumor Samples Yields Insight into Biomarkers of Response and Mechanisms of Resistance to Immune Checkpoint Blockade. Cancer Discov. 6, 827–837 (2016).

44. Nathanson, T. et al. Somatic Mutations and Neoepitope Homology in Melanomas Treated with CTLA-4 Blockade. Cancer Immunol. Res. 5, 84–91 (2017).

45. Friedlander, P. et al. Whole-blood RNA transcript-based models can predict clinical response in two large independent clinical studies of patients with advanced melanoma treated with the checkpoint inhibitor, tremelimumab. J. Immunother. Cancer 5, 67 (2017).

46. Zappasodi, R. et al. CTLA-4 blockade drives loss of Treg stability in glycolysis-low tumours. Nature 591, 652–658 (2021).

47. Weber, J. S. et al. Sequential administration of nivolumab and ipilimumab with a planned switch in patients with advanced melanoma (CheckMate 064): an open-label, randomised, phase 2 trial. Lancet Oncol. 17, 943–955 (2016).

48. Xu, W. & McArthur, G. Cell Cycle Regulation and Melanoma. Curr. Oncol. Rep. 18, 34 (2016).

49. Walker, G. J. et al. Virtually 100% of melanoma cell lines harbor alterations at the DNA level within CDKN2A, CDKN2B, or one of their downstream targets. Genes. Chromosomes Cancer 22, 157–163 (1998).

50. Dasika, G. K. et al. DNA damage-induced cell cycle checkpoints and DNA strand break repair in development and tumorigenesis. Oncogene 18, 7883–7899 (1999).

51. Shain, A. H. et al. Genomic and Transcriptomic Analysis Reveals Incremental Disruption of Key Signaling Pathways during Melanoma Evolution. Cancer Cell 34, 45–55.e4 (2018).

52. Tse, A. K.-W. et al. Sensitization of melanoma cells to alkylating agent-induced DNA damage and cell death via orchestrating oxidative stress and IKKβ inhibition. Redox Biol. 11, 562–576 (2017).

53. Hayes, M. T., Bartley, J. & Parsons, G. In vitro evaluation of fotemustine as a potential agent for limb perfusion in melanoma. Melanoma Res. 8, 67 (1998).

54. Barnaba, N. & LaRocque, J. R. Targeting cell cycle regulation via the G2-M checkpoint for synthetic lethality in melanoma. Cell Cycle 20, 1041–1051 (2021).

55. Chan, P. Y. et al. The synthetic lethal interaction between CDS1 and CDS2 is a vulnerability in uveal melanoma and across multiple tumor types. Nat. Genet. 57, 1672–1683 (2025).

56. Yang, C., Tian, C., Hoffman, T. E., Jacobsen, N. K. & Spencer, S. L. Melanoma subpopulations that rapidly escape MAPK pathway inhibition incur DNA damage and rely on stress signalling. Nat. Commun. 12, 1747 (2021).

57. Deng, Z. et al. TGF-β signaling in health, disease and therapeutics. Signal Transduct. Target. Ther. 9, 61 (2024).

58. Akhurst, R. J. & Derynck, R. TGF-β signaling in cancer – a double-edged sword. Trends Cell Biol. 11, S44–S51 (2001).

59. Cui, W. et al. TGFβ1 Inhibits the Formation of Benign Skin Tumors, but Enhances Progression to Invasive Spindle Carcinomas in Transgenic Mice. Cell 86, 531–542 (1996).

60. Loos, B. et al. TGFβ signaling sensitizes MEKi-resistant human melanoma to targeted therapy-induced apoptosis. Cell Death Dis. 15, 925 (2024).

61. Coffey Jr, R. J., et al. Selective Inhibition of Growth-Related Gene Expression in Murine Keratinocytes by Transforming Growth Factor β. Mol. Cell. Biol. 8, 3088–3093 (1988).

62. Javelaud, D. et al. Stable overexpression of Smad7 in human melanoma cells inhibits their tumorigenicity in vitro and in vivo. Oncogene 24, 7624–7629 (2005).

63. Förstenberger, G. et al. Stimulatory role of transforming growth factors in multistage skin carcinogenesis: Possible explanation for the tumor-inducing effect of wounding in initiated nmri mouse skin. Int. J. Cancer 43, 915–921 (1989).

64. Cui, W., Kemp, C. J., Duffie, E., Balmain, A. & Akhurst, R. J. Lack of Transforming Growth Factor-β1 Expression in Benign Skin Tumors of p53null Mice Is Prognostic for a High Risk of Malignant Conversion1. Cancer Res. 54, 5831–5836 (1994).

65. Glick, A. B. et al. Loss of expression of transforming growth factor beta in skin and skin tumors is associated with hyperproliferation and a high risk for malignant conversion. Proc. Natl. Acad. Sci. 90, 6076–6080 (1993).

66. Arrick, B. A. et al. Altered metabolic and adhesive properties and increased tumorigenesis associated with increased expression of transforming growth factor beta 1. J. Cell Biol. 118, 715–726 (1992).

67. Tuncer, E. et al. SMAD signaling promotes melanoma metastasis independently of phenotype switching. J. Clin. Invest. 129, 2702–2716 (2019).

68. Liu, J. et al. TGF-β in tumor development and progression: mechanisms and therapeutics. Mol. Biomed. 7, 9 (2026).

69. Derynck, R., Turley, S. J. & Akhurst, R. J. TGF-β biology in cancer progression and tumor immunotherapy. Nat. Rev. Clin. Oncol. 18, 9–34 (2021).

70. Chen, B., Mu, C., Zhang, Z., He, X. & Liu, X. The Love-Hate Relationship Between TGF-β Signaling and the Immune System During Development and Tumorigenesis. Front. Immunol. 13, 891268 (2022).

71. Xu, J., Lamouille, S. & Derynck, R. TGF-beta-induced epithelial to mesenchymal transition. Cell Res. 19, 156–172 (2009).

72. Van Den Hout, M. F. C. M., et al. Melanoma Sequentially Suppresses Different DC Subsets in the Sentinel Lymph Node, Affecting Disease Spread and Recurrence. Cancer Immunol. Res. 5, 969–977 (2017).

73. Pekkonen, P. et al. Lymphatic endothelium stimulates melanoma metastasis and invasion via MMP14-dependent Notch3 and β1-integrin activation. eLife 7, e32490 (2018).

74. Das, S. et al. Tumor cell entry into the lymph node is controlled by CCL1 chemokine expressed by lymph node lymphatic sinuses. J. Exp. Med. 210, 1509–1528 (2013).

75. Pedri, D., Karras, P., Landeloos, E., Marine, J.-C. & Rambow, F. Epithelial-to-mesenchymal-like transition events in melanoma. FEBS J. 289, 1352–1368 (2022).

76. Thiery, J. P., Acloque, H., Huang, R. Y. J. & Nieto, M. A. Epithelial-Mesenchymal Transitions in Development and Disease. Cell 139, 871–890 (2009).

77. Kalluri, R. & Weinberg, R. A. The basics of epithelial-mesenchymal transition. J. Clin. Invest. 119, 1420–1428 (2009).

78. Somasundaram, R. et al. Tumor-infiltrating mast cells are associated with resistance to anti-PD-1 therapy. Nat. Commun. 12, 346 (2021).

79. Kaesler, S. et al. Targeting tumor-resident mast cells for effective anti-melanoma immune responses. JCI Insight 4, (2019).

80. Bosisio, F. M. et al. Plasma cells in primary melanoma. Prognostic significance and possible role of IgA. Mod. Pathol. 29, 347–358 (2016).

81. Davis, J. L. et al. Elevated Blood Neutrophil-to-Lymphocyte Ratio: A Readily Available Biomarker Associated with Death Due to Disease in High Risk Non-Metastatic Melanoma. Ann. Surg. Oncol. 24, 1989–1996 (2017).

82. Cohen, J. T., Miner, T. J. & Vezeridis, M. P. Is the Neutrophil-to-Lymphocyte Ratio a Useful Prognostic Indicator in Melanoma Patients? Melanoma Manag. 7, MMT47 (2020).

83. Reticker-Flynn, N. E. & Engleman, E. G. Lymph nodes: at the intersection of cancer treatment and progression. Trends Cell Biol. 33, 1021–1034 (2023).

84. Davis, S. & Meltzer, P. S. GEOquery: a bridge between the Gene Expression Omnibus (GEO) and BioConductor. Bioinformatics 23, 1846–1847 (2007).

85. Huber, W. et al. Orchestrating high-throughput genomic analysis with Bioconductor. Nat. Methods 12, 115–121 (2015).

86. Ritchie, M. E. et al. limma powers differential expression analyses for RNA-sequencing and microarray studies. Nucleic Acids Res. 43, e47 (2015).

87. Schilder, B. & Skene, N. orthogene: an R package for easy mapping of orthologous genes across hundreds of species. (2022).

88. Subramanian, A. et al. Gene set enrichment analysis: A knowledge-based approach for interpreting genome-wide expression profiles. Proc. Natl. Acad. Sci. 102, 15545– 15550 (2005).

89. Korotkevich, G. et al. Fast gene set enrichment analysis. 060012 Preprint at 10.1101/060012 (2021).

90. Wu, T. et al. clusterProfiler 4.0: A universal enrichment tool for interpreting omics data. The Innovation 2, (2021).

91. Kanehisa, M., Goto, S., Sato, Y., Furumichi, M. & Tanabe, M. KEGG for integration and interpretation of large-scale molecular data sets. Nucleic Acids Res. 40, D109– D114 (2012).

92. Liberzon, A. et al. The Molecular Signatures Database Hallmark Gene Set Collection. Cell Syst. 1, 417–425 (2015).

93. Gentles, A. J. et al. A human lung tumor microenvironment interactome identifies clinically relevant cell-type cross-talk. Genome Biol. 21, 107 (2020).

94. Newman, A. M. et al. Determining cell type abundance and expression from bulk tissues with digital cytometry. Nat. Biotechnol. 37, 773–782 (2019).

95. Csardi, G. & Nepusz, T. The igraph software package for complex network research. InterJournal 1695, 1–9.

96. Shannon, P. et al. Cytoscape: A Software Environment for Integrated Models of Biomolecular Interaction Networks. Genome Res. 13, 2498–2504 (2003).

97. Hao, Y. et al. Dictionary learning for integrative, multimodal and scalable single-cell analysis. Nat. Biotechnol. 42, 293–304 (2024).

98. Danaher, P. et al. Insitutype: likelihood-based cell typing for single cell spatial transcriptomics. 2022.10.19.512902 Preprint at 10.1101/2022.10.19.512902 (2022).

99. Ahuja, G. et al. Multi-agent AI enables evidence-based cell annotation in single-cell transcriptomics. 2025.11.06.686964 Preprint at 10.1101/2025.11.06.686964 (2025).

100. Gittleman, J. L. & Kot, M. Adaptation: Statistics and a Null Model for Estimating Phylogenetic Effects. Syst. Biol. 39, 227–241 (1990).

